# Learning and Sleep Have Divergent Effects on Cytosolic and Membrane-Associated Ribosomal mRNA Profiles in Hippocampal Neurons

**DOI:** 10.1101/2020.07.27.221218

**Authors:** James Delorme, Lijing Wang, Varna Kodoth, Yifan Wang, Jingqun Ma, Sha Jiang, Sara J. Aton

## Abstract

The hippocampus plays an essential role in consolidating transient experiences into long-lasting memories. Memory consolidation can be facilitated by post-learning sleep, although the underlying cellular mechanisms are undefined. Here, we addressed this question using a mouse model of hippocampally-mediated, sleep-dependent memory consolidation (contextual fear memory; CFM), which is known to be disrupted by post-learning sleep loss. We used translating ribosome affinity purification (TRAP) to quantify ribosome-associated RNAs in different subcellular compartments (cytosol and membrane) and in different hippocampal cell populations (either whole hippocampus, Camk2a+ excitatory neurons, or highly active neurons expressing phosphorylated ribosomal subunit S6 [pS6+]). Using RNA-seq, we examined how these transcript profiles change as a function of sleep vs. sleep deprivation (SD) and as a function of prior learning (contextual fear conditioning; CFC). Surprisingly, we found that while many mRNAs on cytosolic ribosomes were altered by sleep loss, almost none were altered by learning. Of the few changes in cytosolic ribosomal transcript abundance following CFC, almost all were occluded by subsequent SD. This effect was particularly pronounced in pS6+ neurons with the highest level of neuronal activity following CFC, suggesting SD-induced disruption of post-learning transcript changes in putative “engram” neurons. In striking contrast, far fewer transcripts on membranebound (MB) ribosomes were altered by SD, and many more mRNAs (and lncRNAs) were altered on MB ribosomes as a function of prior learning. For hippocampal neurons, cellular pathways most significantly affected by CFC were involved in structural remodeling. Comparisons of post-CFC transcript profiles between freely-sleeping and SD mice implicated changes in cellular metabolism in Camk2a+ neurons, and increased protein synthesis capacity in pS6+ neurons, as biological processes disrupted by post-learning sleep loss.

## Introduction

The role of sleep in promoting synaptic plasticity and memory storage (consolidation) in the brain is an enduring mystery (Puentes-Mestril and Aton, 2017). For the past two decades, transcriptomic (Cirelli et al., 2004; Mackiewicz et al., 2007; Vecsey et al., 2012) and proteomic (Cirelli et al., 2009; Noya et al., 2019; Poirrier et al., 2008; Ren et al., 2016) profiling of the mammalian brain after sleep vs. experimental sleep deprivation (SD) has provided insights regarding the general functions of sleep for the brain. For example, observed increases in the abundance of transcripts for immediate early genes and synaptic proteins after SD were the initial basis for the synaptic homeostasis hypothesis for sleep function (Cirelli et al., 2004). The hypothesis proposes that synapses are broadly “downscaled” during sleep. However, the function of such a process in memory consolidation, and it’s occurrence during post-learning sleep, are a matter of debate (Havekes and Aton, 2020). On the other hand, observations from *in vivo* electrophysiology in the sleeping brain has led to conclusion that specific patterns of activity present during learning experiences may be replayed during subsequent sleep. Such a mechanism would be instructive with regard to memory storage (i.e., selectively affecting only highly active “engram neurons” engaged during prior experience and their postsynaptic partners). However, determining whether replay is necessary for memory consolidation has been difficult (Puentes-Mestril and Aton, 2017; Puentes-Mestril et al., 2019), and it is unknown how this process (and other features of brain physiology associated with sleep) would affect intracellular pathways (e.g., those involved in synaptic plasticity).

More recently, transcriptomic and proteomic profiling of synaptic or axonal organelles has been used to better understand the effects of learning (Ostroff et al., 2019) or of sleep vs. wake (Noya et al., 2019) on synaptic function. However, to date, there has been no experimental work aimed at characterizing cellular changes during sleep-dependent memory consolidation - i.e., those occurring as a function of post-learning sleep. Here, we use a well-established mouse model of sleep-dependent memory consolidation - contextual fear memory (CFM) - to study this process. CFM can be encoded in a single learning trial (contextual fear conditioning; CFC), and is consolidated via hippocampus-dependent mechanisms over the next few hours. Critically, CFM consolidation can be disrupted by sleep deprivation (SD) within the first 5-6 h following CFC (Graves et al., 2003; Ognjanovski et al., 2018). Over this same post-CFC time interval, disruption of either neuronal activity (Daumas et al., 2005), transcription (Igaz et al., 2002; Pereira et al., 2019), or translation (Gafford et al., 2011; Tudor et al., 2016) in the dorsal hippocampus can likewise disrupt consolidation. This suggests that an activity- and sleep-dependent mechanism, impinging on biosynthetic pathways in the hippocampus, is essential for consolidation.

To shed light on this putative mechanism, we characterized changes to ribosome-associated mRNAs from different hippocampal cell populations (including Camk2a+ excitatory neurons and highly active neurons expressing phosphorylated S6 [pS6+]) as a function of both sleep vs. SD and prior CFC. By quantifying mRNA profiles on ribosomes differentially localized to the cytosol and cellular membranes, we find that while the majority of changes to transcripts on cytosolic ribosomes vary as a function of sleep vs. wake, the majority of transcript changes on membrane-bound (MB) ribosomes vary as a function of learning. Our findings reveal new subcellular functions for post-learning sleep, and suggest new cellular mechanisms by which sleep could selectively promote memory storage.

## Results

### TRAP-Mediated Isolation of mRNAs From Hippocampal Neuron Subpopulations

To quantify the effects of sleep and learning on hippocampal mRNA translation, we employed two translating ribosome affinity purification (TRAP) techniques. First, to quantify ribosome-associated mRNAs in excitatory neurons, B6.Cg-Tg(Camk2a-cre)T29-1Stl/J mice were crossed to the *B6N.129-Rpl22^tm1.1Psam/J^* mouse line (Sanz et al., 2019; Sanz et al., 2009). Offspring from this cross express hemmatagluttin (HA)-tagged ribosomal protein 22 (HA-Rpl22) in excitatory (Camk2a+) neurons (**Figure 1A, left**). Second, to quantify mRNAs associated with ribosomes in active hippocampal neurons, we used an antibody targeting the terminal phosphorylation sites (Ser244/247) of ribosomal protein S6 (pS6) (Knight et al., 2012) (**Figure 1A, left**). These sites are phosphorylated in neurons as the result of high neuronal activity, by mTOR-dependent kinase S6K1/2 (Biever et al., 2015). This strategy allowed us to compare mRNAs expressed in the whole hippocampus (Input) with those associated with ribosomes in either Camk2a+ or highly active (pS6+) neuronal populations from the same hippocampal tissue. To further test how mRNA translation varies as a function of ribosomes’ subcellular localization, we centrifuged our homogenized hippocampal tissue and collected samples from supernatant (presumptive cytosolic) and pellet (presumptive membrane-containing) fractions (Kratz et al., 2014). From both fractions, we compared whole-hippocampus (Input) transcripts with transcripts isolated by TRAP from excitatory (Camk2a+) and highly active (pS6+) neuron populations (**Figure 1A**).

**Figure 1.**
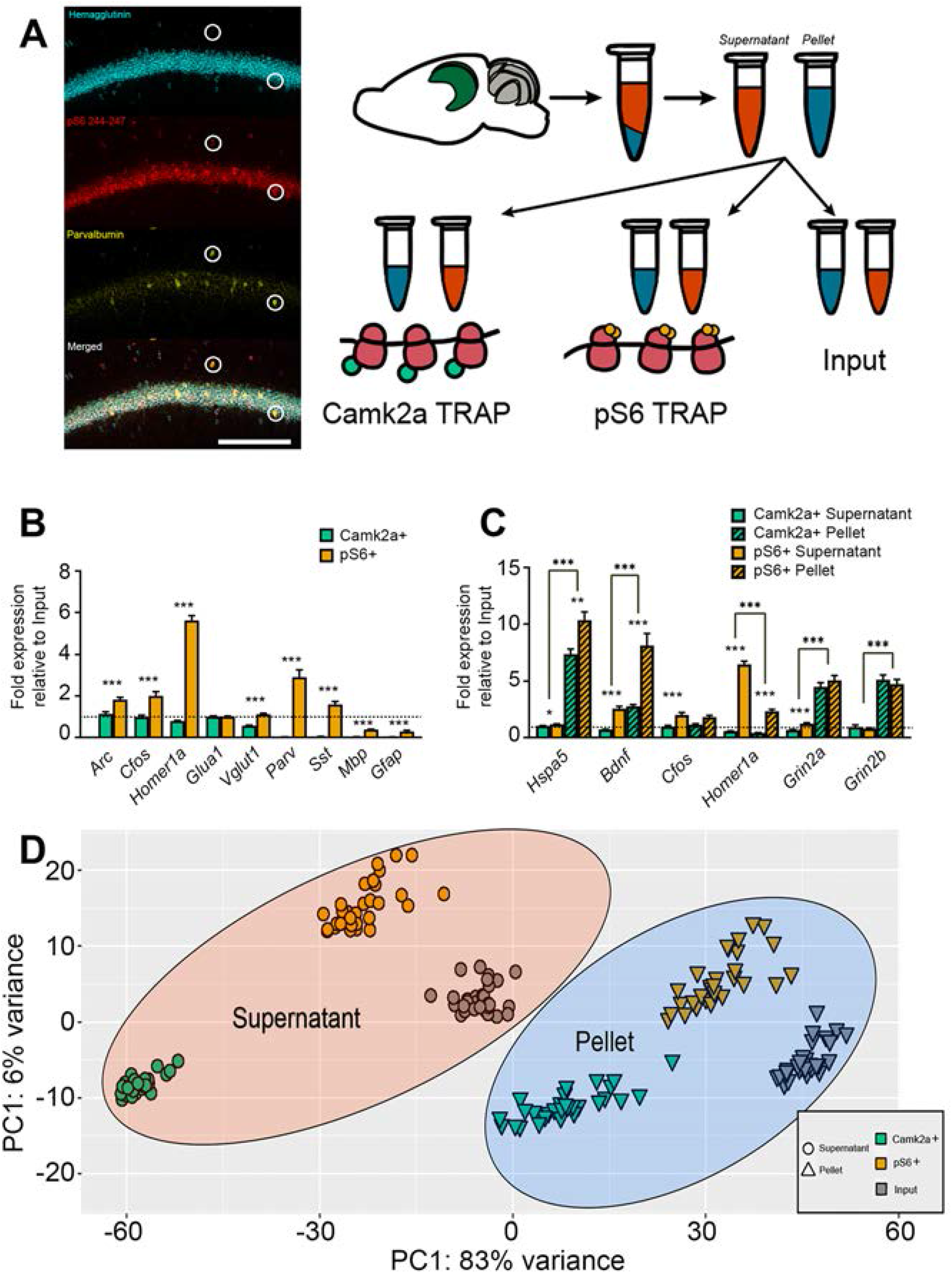
TRAP-based profiling of hippocampal cell populations and isolation of subcellular fractions. **(A) *Left:*** Confocal images showing expression of hemagglutinin (HA, Camk2a), phosphorylated S6 (pS6), and parvalbumin in area CA1 of dorsal hippocampus. Highlighted neurons are parvalbumin+, pS6+, and HA-. Scale bar = 100 μm. ***Right:*** Schematic of protocol for isolating mRNAs from subcellular fractions and different cell populations using TRAP. **(B)** Camk2a+ (cyan) and pS6+ (orange) TRAP mRNA enrichment values were calculated (vs. Input) for activity-dependent (*Arc*, *Cfos, Homer1a*), excitatory neuron (*Glua1, Vglut1*), inhibitory neuron (*Parv, Sst*), and glial (*Mbp, Gfap*) transcripts. *** indicates *p* < 0.001 for enrichment value differences between Camk2a+ and pS6+ neuronal populations (Student’s t-test, *n* = 7/group). **(C)** Camk2a+ and pS6+ TRAP enrichment in supernatant (solid bars) and pellet (hatched bars) fractions (vs. Input) for transcripts encoding secreted (*Bdnf*), transmembrane (*Grin2a, Grin2b*), endoplasmic reticulum (*Hspa5*), and cytosolic (*Cfos, Homer1a*) proteins. (Student’s t-test, *n* = 9/group, *, **, and *** indicate *p* < 0.05, *p* < 0.01, and *p* < 0.001, respectively). **(D)** PCA plot (VST, Deseq2) for RNAseq data (from *n* = 30 hippocampal samples) from the three cell populations (Input, Camk2a+ neurons, and pS6+ neurons) and two fractions (supernatant and pellet).

Using quantitative PCR (qPCR), we first validated cell type-specific gene expression from Camk2a+ and pS6+ cell populations. Relative to Input mRNA, Camk2a+ mRNA displayed similar levels of *Arc, Cfos, Homer1a, Glua1*, and *Vglut1*, with reduced expression of interneuron and glial cell markers. Compared to Input, highly activated pS6+ neurons’ mRNA profiles displayed significant enrichment in activity-dependent transcripts (*Arc, Cfos, Homer1a*) and interneuron-specific transcripts (*Pvalb, Sst*), and comparable levels of excitatory neuron-specific mRNAs (*Glua1, Vglut1*) (**Figure 1A,B**).

We next used qPCR for preliminary validation of sub-cellular enrichment of mRNAs expressed in supernatant and pellet fractions. Previous reports have found that isolating pellet ribosomes enriches for endoplasmic reticulum (ER) and dendritic localized ribosomes (Kratz et al., 2014). To test whether fractions differently enriched genes trafficked to the ER and dendrites we first analyzed *Hspa5.* Encoding the resident ER chaperone BIP, *Hspa5* was significantly more enriched in the pellet fractions than the cytosolic fractions of both Camk2a+ and pS6+ neurons (**Figure 1C**; Camk2a+: supernatant - 1.03 × Input, pellet - 7.38 × Input, *p* < 0.001, Student’s t-test; pS6+: supernatant - 1.17 × Input, pellet - 10.33 × Input, *p* < 0.001). Similarly, *Bdnf, Grin2a*, and *Grin2b* mRNAs (encoding the secreted growth factor and glutamatergic receptor subunits) were more enriched on ribosomes isolated from the pellet fraction compared to the supernatant fraction. In contrast, *Homer1a* (encoding the truncated version of the synaptic scaffolding protein Homer1, present in cytosol) was more abundant on supernatant ribosomes in both neuron populations, and *Cfos* was equally abundant in both fractions. These results suggest that ribosome-associated transcripts observed in pellet and supernatant fractions encode proteins with predicted enrichment on cell membranes and in cytosol, respectively. We further characterized mRNAs taken from different cell populations (Camk2a+, pS6+, Input) and subcellular fractions (supernatant, pellet) using a non-biased approach - RNA-seq. PCA analysis of the full RNA-seq data set revealed six discrete clusters of mRNA expression profiles, based on the origin of the samples (**Figure 1D**).

### Supernatant and Pellet Fractions Distinguish Transcripts Localized to Cytosolic and Membrane-Bound (MB) Ribosomes

To characterize transcripts differentially localized to the supernatant or pellet fraction, we next calculated the relative mRNA abundance between the two fractions from Camk2a+ neurons (**Figure 2A**), pS6+ neurons (**Figure 3A**), and Input (i.e., whole hippocampus, **Figure S1A**) using Deseq2 (Love et al., 2014). The top 2000 most differentially expressed transcripts between the two fractions (based on adjusted *p* value) were then characterized using DAVID’s cellular component annotation (Dennis Jr. et al., 2003). Confirming our initial validation (**Figure 1B**), supernatant-enriched transcripts from both neuron populations (and Input) encoded proteins with functions localized to the cytoplasm and nucleus. Pellet-enriched transcripts encoded proteins with functions localized to the plasma membrane, endoplasmic reticulum, Golgi apparatus, and synapses (**Figure 2B, Figure 3B, Figure S1B**). Thus for subsequent analyses, we refer to supernatant and pellet fractions as cytosolic and membrane-bound (MB), respectively. Signaling and metabolic pathways enriched among cytosolic ribosome-associated mRNAs were assessed using Ingenuity Pathway Analysis [IPA] canonical pathways (**Figure 2C-D, Figure 3C-D**). Here, we identified cytosol-localized cellular pathways including ubiquitination, nucleotide excision repair, hypoxia signaling, and sumoylation pathways. In contrast, MB fraction-enriched transcripts represented signaling pathways involved in synaptic (GABAergic receptor, glutamatergic receptor, and endocannabinoid signaling) and endoplasmic reticulum (e.g., unfolded protein response) functions (**Figure 2C-D, Figure 3C-D**).

**Figure 2.**
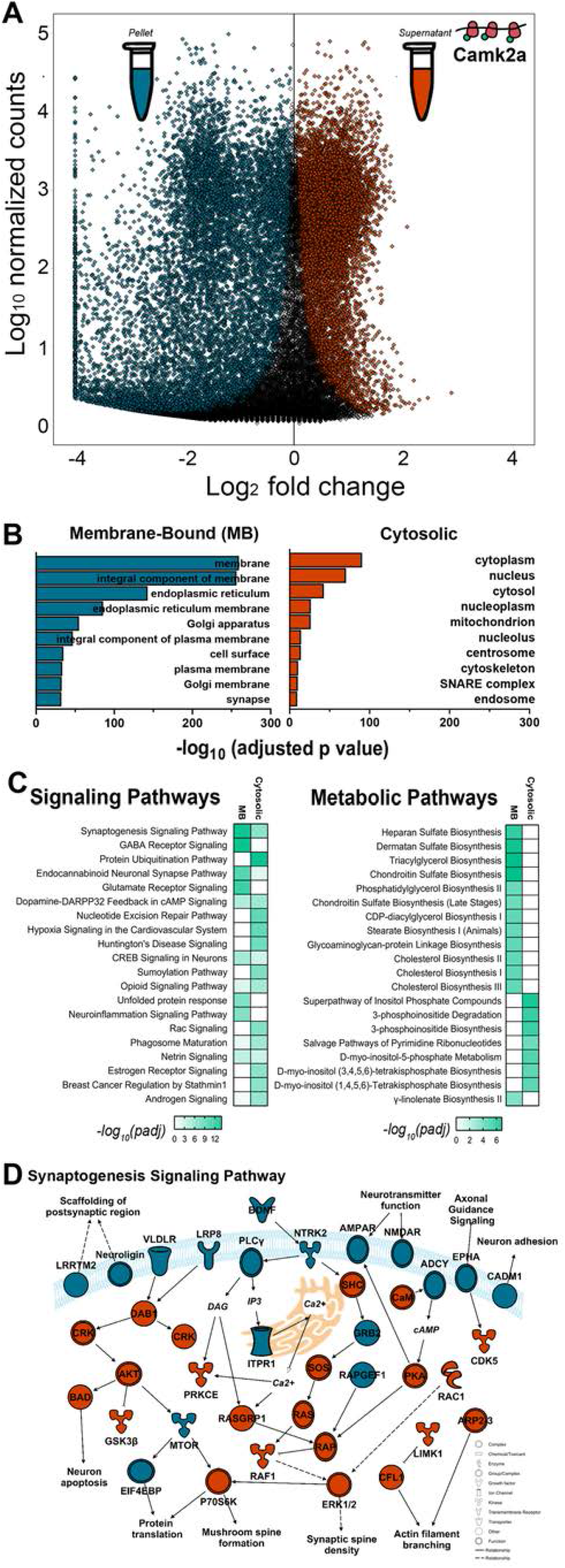
Cytosolic and membrane protein-encoding transcripts in Camk2a+ neurons are preferentially enriched in supernatant and pellet ribosomal fractions respectively. **(A)** Volcano plot of transcripts significantly enriched in pellet (red) and supernatant (blue) cell fractions of Camk2a+ neurons. Of the 28,071 transcripts detected, 7,651 (27%) were significantly (*p_adj_* < 0.1) enriched in the supernatant (cytosolic) fraction, and 10,911 (39%) were significantly enriched in the pellet (MB) fraction. Complete transcript list in **Supplemental Table S1**. **(B)** Top 10 cellular component localizations (from DAVID) of the 2000 transcripts most significantly enriched (based on adjusted *p* value) in either pellet (MB) or supernatant (cytosolic) fractions. **(C)** Top 20 most-enriched signaling and metabolic pathways represented by the 2000 most-enriched transcripts in Camk2a+ cytosolic or MB fractions. **(D)** Illustration of the synaptogenesis signaling pathway (IPA) with proteins shaded by their respective transcripts’ preferential localization in cytosolic (blue) and MB (red) fractions. Functional category analysis available in **Supplemental Table S1.**

**Figure 3.**
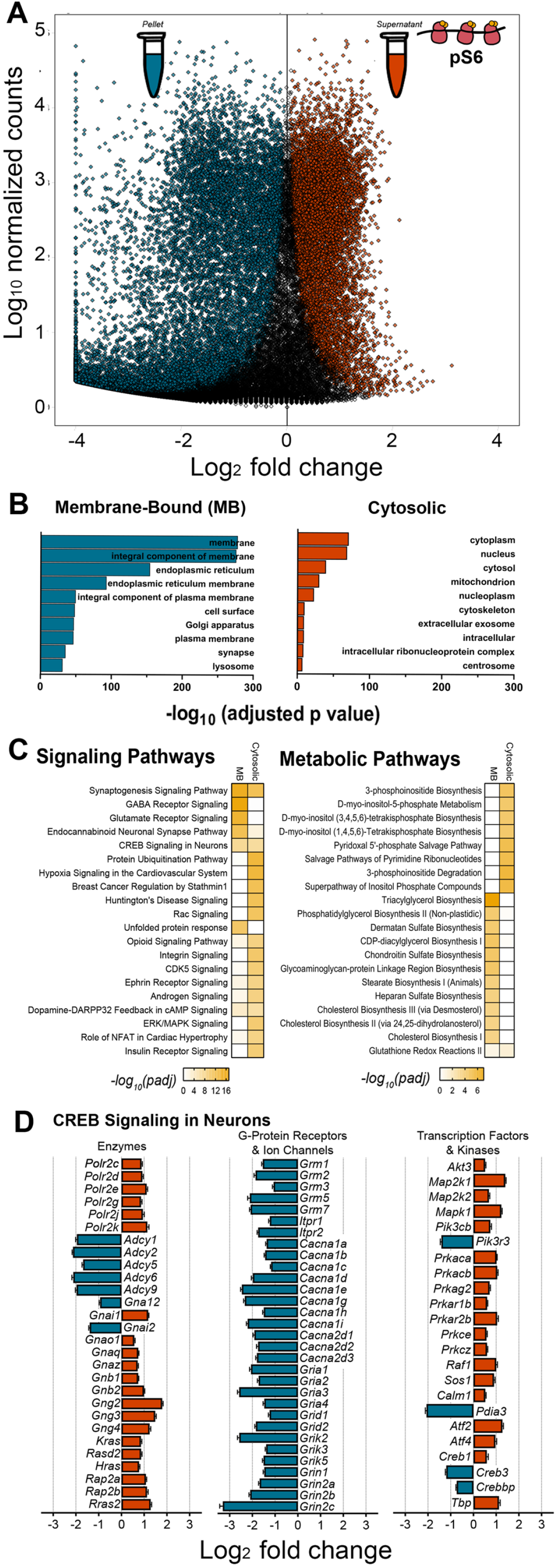
Differential expression of mRNAs encoding intracellular signaling pathway components in cytosolic vs. MB ribosomal fractions from highly active (pS6+) neurons. **(A)** Volcano plot of transcripts significantly enriched in pellet (red) and supernatant (blue) cell fractions. Of the 34,657 transcripts detected, 8,030 (23%) were significantly (*p_adj_* < 0.1) enriched in the supernatant (cytosolic) fraction, and 14,244 (41%) were significantly enriched in the pellet (MB) fraction. Complete transcript list in **Supplemental Table S2**. **(B)** Top 10 cellular component localizations (from DAVID) of the 2000 transcripts which were most significantly enriched (based on adjusted *p* value) in either pellet (MB) or supernatant (cytosolic) fractions of pS6+ neurons. **(C)** Top 20 most-enriched signaling and metabolic pathways represented by the 2000 most-enriched transcripts in pS6+ cytosolic or MB fractions. **(D)** Log2FC values indicating enrichment of transcripts from the Creb1 signaling pathway in the cytosolic (red) or MB (blue) fractions (subcategorized by encoded protein type). Functional category analysis available in **Supplemental Table S2**

To further investigate subcellular localization of transcripts representing cellular pathways critically involved in hippocampal function, we examined signaling pathways enriched in both cytosolic and MB fractions. Because both learning and sleep affect synaptic structure and function (Bruning et al., 2019; Noya et al., 2019; Raven et al., 2019; Spano et al., 2019), we first focused on the synaptogenesis signaling pathway. In Camk2a+ neurons, MB-enriched transcripts encoded secreted proteins (e.g., *Bdnf*), transmembrane proteins including AMPA, NMDA, and ephrin receptors (e.g., *Gria1, Gria2, Gria3, Grin2a, Grin2b, Grin2c, Epha1, Epha2*) and membrane-associated enzymes (*Plcγ*) (**Figure 2D**). Cytosol-enriched mRNAs encoded intracellular complexes including adaptor proteins (*Crk, Shc*) and kinases (*Cdk5, Lmk1, Gsk3b, Mapk1, Mapk2, P70S6K*) in the synaptogenesis pathway (**Figure 2D**). Components of the CREB signaling pathway (another known target of both learning and sleep) (Abel et al., 1997; Kandel, 2012; Luo et al., 2013; Vecsey et al., 2009; Vecsey et al., 2007) was also differentially enriched on cytosolic vs. MB ribosomes, in both Camk2a+ and pS6+ neuronal populations (**Figure 2C, Figure 3C**). mRNAs encoding enzymes in the CREB pathway were selectively localized to ribosomes in either compartment of neuronal populations (e.g., *Polr2c*, encoding the RNA polymerase subunit, in the cytosolic fraction; *Adcy1*, encoding adenylate cyclase, in the MB fraction). mRNAs encoding G-protein coupled receptors and ion channels (metabotropic glutamate receptors, endoplasmic reticulum IP3 receptors, calcium channel subunits, AMPA and NMDA receptor subunits) localized exclusively to the MB fraction, while those encoding transcription factors and kinases localized primarily to the cytosolic fraction (**Figure 3D**). In contrast to Camk2a+ and pS6+ neuron populations, the CREB signaling pathway was not represented among mRNAs differentially localized to subcellular fractions of Input (i.e., whole hippocampus; **Figure S1**), suggesting that differential localization of mRNAs encoding CREB signaling components are more pronounced in neurons than other hippocampal cell types.

Overall, functional categories represented in the two subcellular fractions in Input RNA followed a pattern similar to that seen in Camk2a+ and pS6+ neuronal populations. However, in contrast to ribosome-associated transcript profiles from neuronal populations, the signaling pathway category that was most represented by mRNAs differentially localized between the two Input fractions was the protein ubiquitination pathway (**Figure S1**). This may indicate more dramatic subcellular segregation of mRNAs encoding ubiquitin pathway components in nonneuronal hippocampal cell types (i.e., glial cells).

### Learning and Sleep Loss Have Divergent Effects on Cytosolic and MB Ribosome-Associated mRNA Profiles

Because both learning and sleep alter hippocampal activity, intracellular signaling, and function (Havekes et al., 2016; Ognjanovski et al., 2018; Ognjanovski et al., 2014; Vecsey et al., 2009), we next tested how cytosolic and MB ribosome-associated transcripts in different neuron types were affected by prior training on a hippocampus-dependent memory task (contextual fear conditioning [CFC]). We also tested how these transcript profiles were affected by a brief (3-h) period of subsequent sleep or sleep deprivation (which is sufficient to disrupt contextual fear memory (CFM) consolidation) (Graves et al., 2003; Ognjanovski et al., 2018; Vecsey et al., 2009). At lights on (i.e., the beginning of the rest phase), mice were either left in their home cage (HC) or underwent single-trial CFC (placement in a novel chamber followed by delivery of a foot shock). Over the next 3 h, mice in CFC and HC control groups were either permitted *ad lib* sleep (Sleep) or were sleep deprived (SD) in their home cage by gentle handling (**Figure 4A**). These manipulations were followed by RNA isolation and sequencing as described above. Effects of learning and sleep loss on mRNA abundance were quantified for each cell population (Camk2a+, pS6+, or Input) and subcellular fraction (cytosolic or MB; e.g., pS6+ MB) to preserve gene-level inferences made by the Deseq2 model.

**Figure 4.**
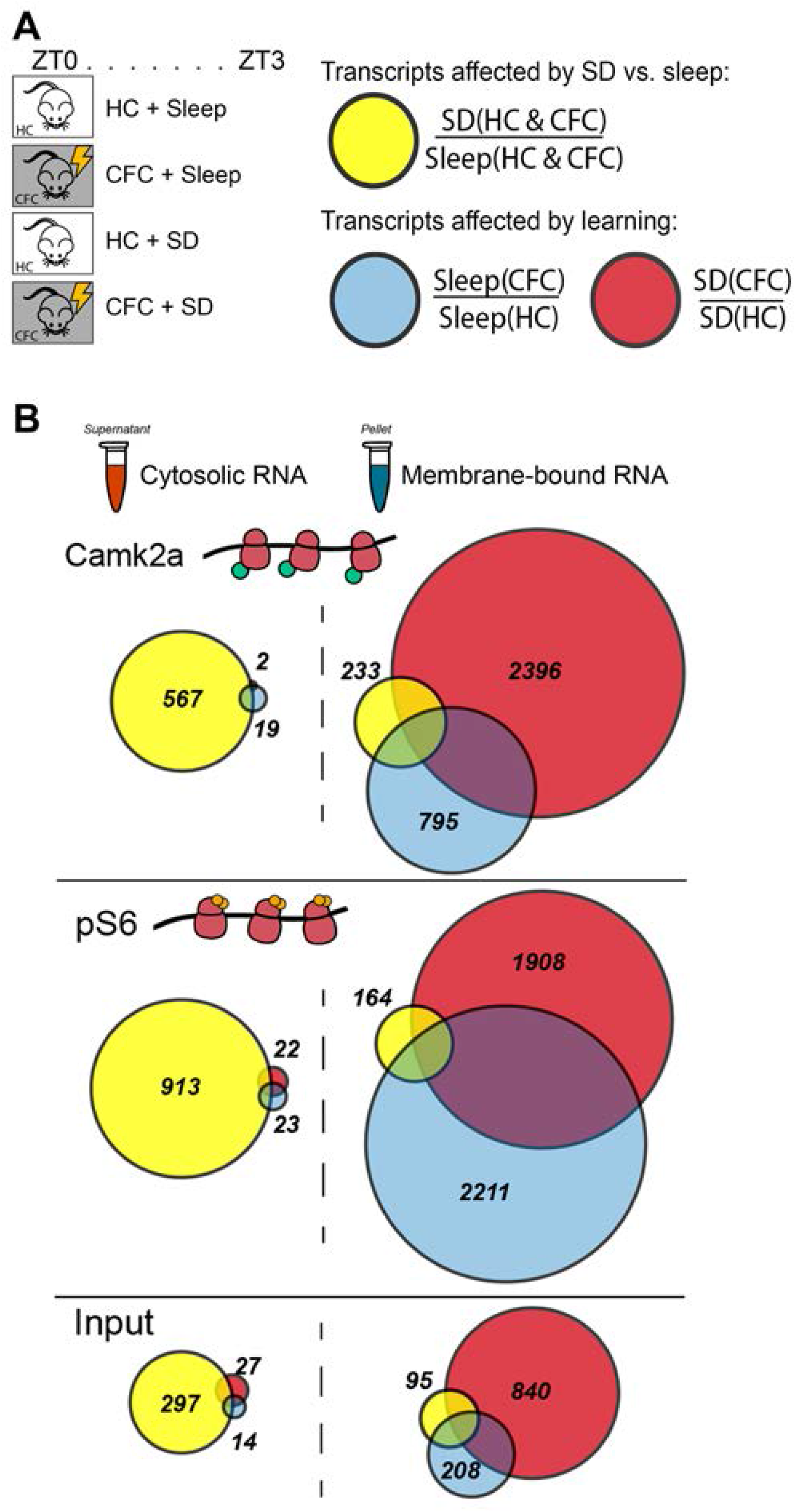
Cytosolic ribosomal transcripts are altered primarily by SD, while MB ribosomal transcripts are altered primarily by learning. **(A) *Left:*** Experimental paradigm for RNA-seq experiments. At lights on, mice were either left in their home cage (HC) or underwent single-trial CFC. All mice were then either permitted *ad lib* sleep or were sleep deprived (SD) over the following 3 h. ***Right:*** Transcript comparisons for quantifying effects of SD (yellow) included both HC and CFC animals. To quantify effects of CFC, CFC + Sleep (Blue) and CFC + SD (Red) mice were analyzed separately. Following behavioral manipulations, cytosolic and MB fractions for different cell populations were isolated as described in **Figure 1. (B)** Proportional Venn diagrams reflect the number of significantly altered transcripts in each cell populations and subcellular fractions (i.e., Camk2a+/Membrane-Bound), based on comparisons shown in **A**. Complete transcript lists for each comparison are available in **Supplemental Table 4-6.**

We first assessed the specific effects of sleep deprivation alone (comparing SD and Sleep conditions) by combining data sets from naive (HC) and recently trained (CFC) mice (**Figure 4A, Yellow**). We then quantified the effects of learning (comparing CFC and HC conditions) separately in Sleeping and SD mice (**Figure 4A, Red[SD], Blue[Sleep]**). Venn diagrams (shown in **Figure 4B**) show the proportional changes in mRNAs resulting from these comparisons. SD had a relatively large effect on cytosolic ribosomal mRNAs (Camk2a+: 567 transcripts, pS6+: 913 transcripts, Input: 297 transcripts) compared to the effect of learning, which had extremely modest effects on cytosolic ribosomal transcripts. Conversely, MB ribosomal mRNAs were dramatically altered by learning in both SD (CFC + SD - Camk2a+: 2,396 transcripts, pS6+: 1,908 transcripts, Input: 840 transcripts) and Sleep groups (CFC + Sleep - Camk2a+: 795 transcripts, pS6+: 2,211 transcripts, Input: 208 transcripts). In contrast, relatively few MB ribosomal mRNAs were altered by SD (Camk2a+: 233 transcripts, pS6+: 164 transcripts, Input: 95 transcripts). These results suggest that SD and learning differentially affect ribosomal mRNA profiles based on their subcellular localization - with SD (vs. Sleep) having more pronounced effects in the cytosol, and learning having more pronounced effects on MB ribosomes. Relatively few transcripts affected by sleep or learning overlapped between different cell populations, as shown in **Figure 5**.

**Figure 5.**
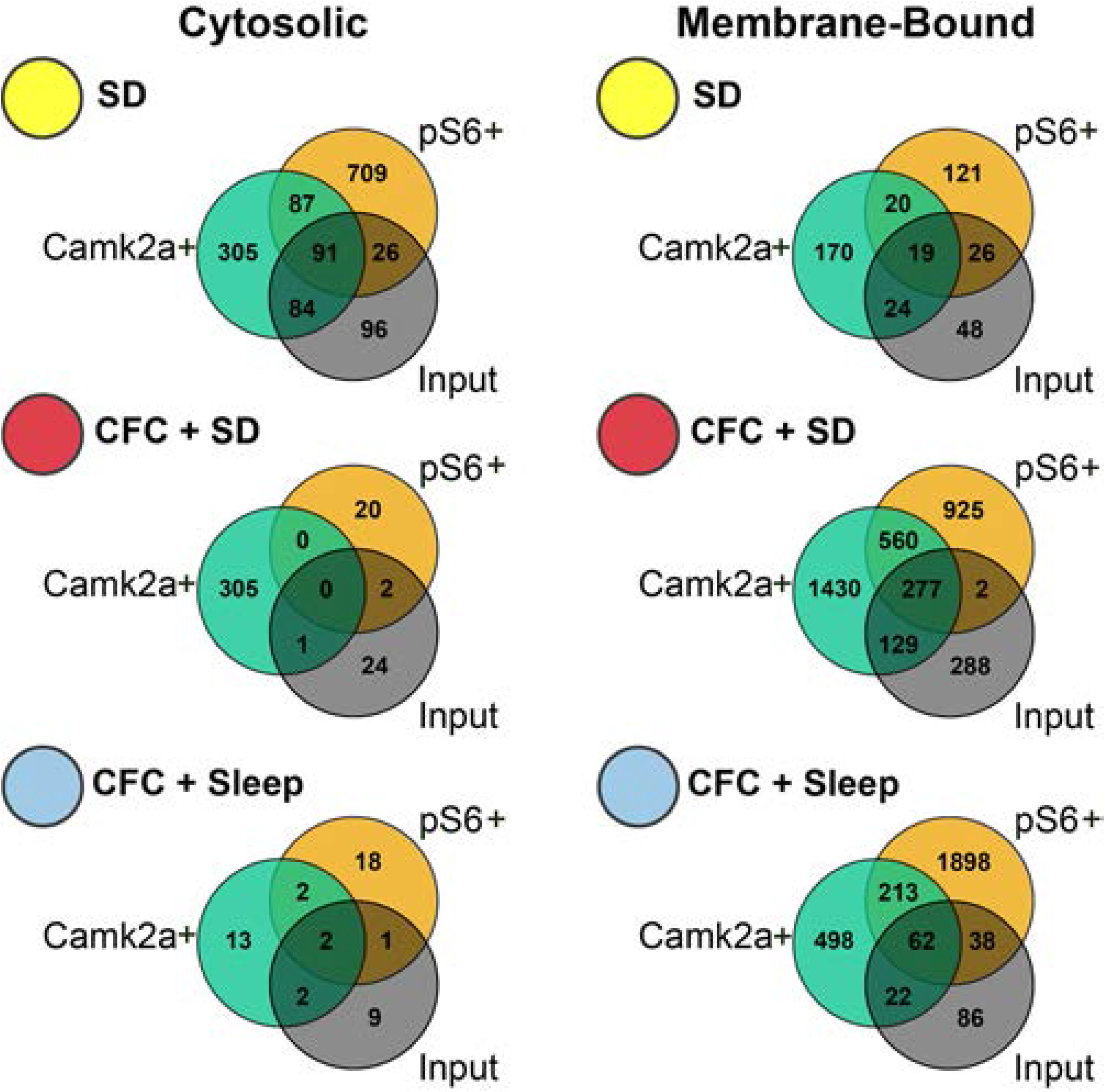
Overlap of transcripts affected by SD or learning in different cell populations. Transcripts altered by SD and CFC in different subcellular fractions in Camk2a+, pS6+, and Input populations were used to construct Venn diagrams. Full transcript lists are included in **Supplemental Table 7**.

### SD Primarily Affects Cytosolic Ribosomal mRNAs Involved in Transcription Regulation

Significantly more cytosolic ribosome-associated mRNAs were altered by SD (compared with relatively few changes driven by CFC) in Camk2a+ neurons (SD: 567 transcripts, CFC: 20 transcripts), pS6+ neurons (SD: 913 transcripts, CFC: 43 transcripts), and to a lesser extent, Input (whole hippocampus) mRNA (SD: 297 transcripts, CFC: 37 transcripts) (**Figure 6**). Therefore, we sought to characterize the molecular and cellular pathways altered by SD-induced changes to cytosolic ribosome transcripts (**Figure 6A**). Molecular functions most affected by SD alone (based on adjusted *p* values <0.1 for transcripts using IPA annotation) overwhelmingly favored transcriptional regulation and RNA processing in both Camk2a+ and pS6+ neurons, as well as in Input (**Figure 6B, Top**). Previous transcriptome analysis has shown that mRNAs encoding transcription regulators are more abundant following brief SD in the hippocampus (*Fos, Elk1, Nr4a1, Creb, Crem1*) (Vecsey et al., 2012) and cortex (*Per2, Egr1, Nr4a1*) (Cirelli et al., 2004). Consistent with those findings, SD increased the abundance of multiple mRNAs encoding transcription factors and upstream regulators in all cell populations, including *E2f6, Elk1, Erf, Fosl1, Fosl2, Fos, Fosb, Lmo4, Taf12, Xbp1, Atf7, Artnl2, Atoh8, Bhlhe40(Dec1), Crebl2, Crem, Egr2, Nfil3*, and *Ubp1.* Transcripts affected in Camk2a+ and pS6+ neurons overlapped partially with those reported in previous SD experiments, including components of pathways for AMPK, PDGF, ERK/MAPK, IGF-1 and endoplasmic reticulum stress signaling (**Figure 6B, Bottom**) (Naidoo et al., 2005; Tudor et al., 2016; Vecsey et al., 2012). However, only a small fraction (12-16%) of the 511 mRNAs previously reported to be altered by SD in whole hippocampus (Vecsey et al., 2012) overlapped with SD-affected mRNAs in cytosolic fractions of any cell population (**Figure S2**). In pS6+ neurons only, mRNAs encoding components of the PI3K/AKT and TGF-B signaling pathways were downregulated in the cytosolic fraction after SD (**Figure 6B, Bottom**), suggesting that activity in these pathways may be higher during sleep.

**Figure 6.**
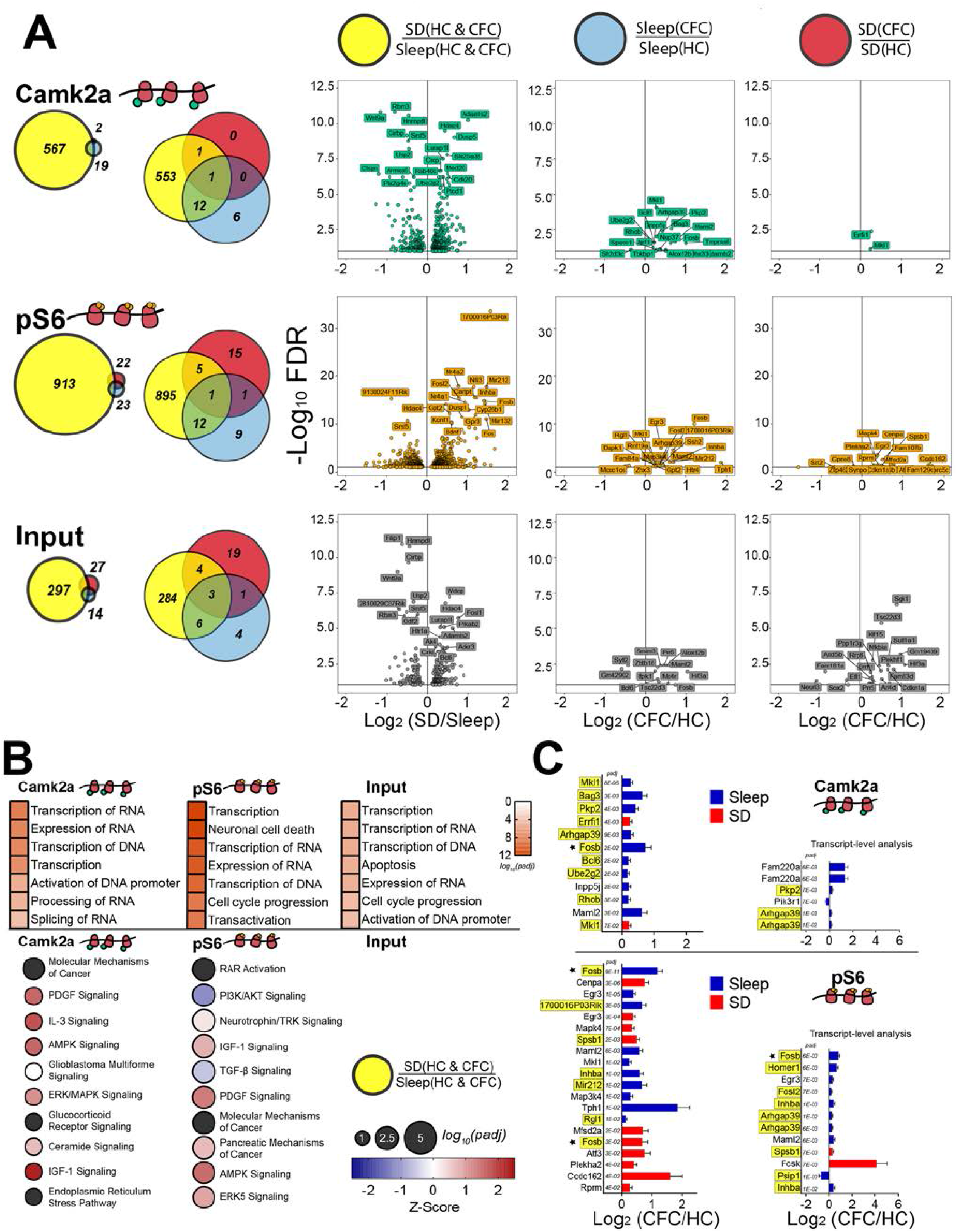
mRNAs altered by SD on cytosolic ribosomes encode transcriptional regulators. **(A) *Left:*** Proportional and overlapping Venn diagrams of transcripts significantly altered by SD, CFC + Sleep, and CFC + SD in cytosolic fractions from Camk2a+ neurons, pS6+ neurons, and Input. ***Right:*** Volcano plots of Deseq2 results for transcripts measured in each condition. **(B) *Top*:** The 7 most-enriched molecular and cellular function categories (ranked by *p*_adj_ value) for transcripts altered by SD alone in Camk2a+ neurons, pS6+ neurons, and Input. ***Bottom:*** The 10 most-enriched canonical pathways of SD-affected transcripts are listed in order *p*_adj_ value (indicated by circle diameter), with z-scores indicating direction of pathway regulation (indicated by hue). There were no significant canonical pathways present in the Input fraction. **(C) *Left:*** The 10 transcripts most significantly affected in CFC + Sleep (blue) and CFC + SD (red) conditions for Camk2a+ (***top***) and pS6+ (***bottom***) neurons, ranked by *p*_adj_ value. Transcripts that were also significantly altered as a function of SD alone are highlighted in yellow. ***Right:*** Results of transcript-level analysis (Yi et al., 2018), show transcripts for transcript isoforms altered in Camk2a+ (***top***) and pS6+ (***bottom***) neurons following CFC. Transcript isoforms that were significantly altered as a function of SD are highlighted in yellow. Functional category analysis available in **Supplemental Table S8.**

We next performed upstream regulator analysis to characterize transcript changes due to SD-associated transcriptional regulation. Results of this analysis provide both a *p*-value for the significance of mRNAs’ regulation by a specific common upstream regulator, and a z-score indicating the direction of the regulated mRNAs’ fold change (i.e., transcriptional activation or suppression). Taken together these values predict the activation state of specific gene regulator complexes during SD (Kramer et al., 2014). In line with prior meta-analysis of SD-induced transcripts (Wang et al., 2010), Creb1 was identified as the transcriptional regulator whose downstream targets’ were most consistently affected across all cytosolic (**Figure 7A**). *Creb1* transcript itself was not increased following SD, although *Crebl2* and *Crem* mRNAs were both increased, and multiple Creb1 transcriptional targets (e.g., *Fos, Arc, FosB, Egr2, Nfil3, Nr4a2, Bag3, Irs2*) were upregulated in the cytosolic fraction of both Camk2a+ and pS6+ neuronal populations (**Figure 7B, Figure S3**) following SD.

**Figure 7.**
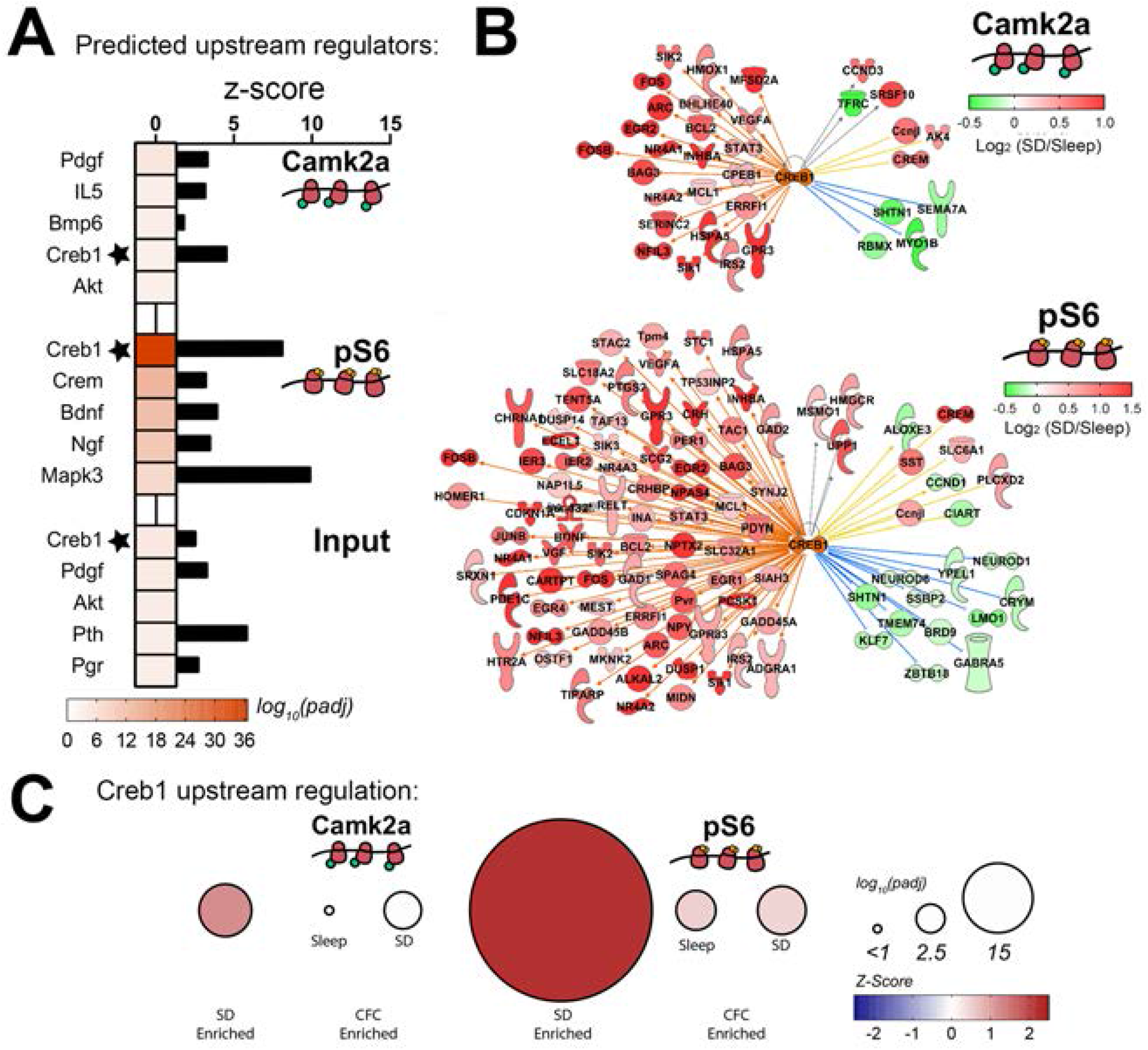
Creb1 target transcripts are upregulated on cytosolic ribosomes after SD. **(A)** Z-scores for the 5 upstream transcriptional regulators whose target transcripts were most significantly affected by SD, ranked by *p*_adj_ values. **(B)** Networks of Creb1 transcriptional targets altered by SD in Camk2a+ (***top***) and pS6+ (***bottom***) neurons. Color of arrows from Creb1 to transcripts indicates the predicted direction of transcriptional regulation - orange (with red transcript symbols) denotes transcripts predicted to be upregulated by Creb1 which are upregulated following SD; blue (with green transcript symbols) indicates transcripts predicted to be repressed by Creb1 which are repressed by SD; yellow indicates SD-related changes that do not match predicted regulation by Creb1; grey indicates undermined effects of Creb1 on transcript levels. **(C)** Relative Creb1 network regulation *p*_adj_ values and z-scores are plotted for transcripts altered on cytosolic ribosomes by SD, CFC + Sleep, and CFC + SD in Camk2a+ and pS6+ cell populations. Upstream regulator analysis results are included in **Supplemental Table S9.** Cytosolic input CREB network analyses are available in **Supplemental Figure 3**.

### CFC-Induced Changes in Cytosolic Ribosome-Associated mRNAs Are Occluded by Subsequent SD

Hippocampal fear memory consolidation in the hours following CFC relies on both sleep (Graves et al., 2003; Ognjanovski et al., 2018; Vecsey et al., 2009) and CREB-mediated transcription (Katche et al., 2010; Rao-Ruiz et al., 2019). With Creb1 activity high during SD, we were curious what effect SD would have on the abundance of ribosome-associated transcripts involved in memory consolidation. As discussed above, few cytosolic ribosome-associated mRNAs were altered by CFC (**Figure 4, 6**). In Camk2a+ neurons, 19 cytosolic ribosome-associated transcripts were altered (compared to HC controls) in CFC + Sleep mice, whereas only 2 transcripts were altered in CFC + SD mice. Of those, most (13/19 from CFC + Sleep, 2/2 from CFC + SD) were also increased by SD alone (**Figure 6A**). Ribosome-associated mRNAs affected by both SD and CFC in Camk2a+ neurons included activity-dependent transcripts such as *Fosb, Arhgap39(Vilse*), and *Errfi1* (**Figure 6C, Yellow**). For comparison, for Input (i.e., whole hippocampus), slightly more cytosolic ribosome-associated mRNAs were altered after CFC + SD (27 transcripts) vs. CFC + Sleep (14 transcripts). Of these, 6/27 and 9/14, respectively, were similarly affected by SD alone (**Figure 6A**).

These initial data suggested that changes in cytosolic ribosome-associated transcripts following SD could occlude transcript changes triggered by CFC. To better characterize how this might affect neurons most activated by CFC (which could represent CFC “engram neurons”), we compared cytosolic transcripts affected by CFC and SD in pS6+ neurons. While similar numbers of transcripts were altered by CFC followed by sleep or SD (23 vs. 22), only two transcripts (*Fosb* and *Egr3*) were similarly affected in both CFC + Sleep and CFC + SD mice (**Figure 6A, C**). Of the transcripts altered on cytosolic ribosomes after CFC in pS6+ neurons, several (12/23 of those affected in freely sleeping mice, 6/22 of those affected in SD mice) were also regulated by SD (**Figure 6A)**, including *Fosb, Egr3, Arghap39(Vilse), Gpr3, Ssh2, Inhba, Rnf19a*, and *Cdkn1a* (**Figure 6C, Yellow**). Because SD alters a significant number of transcripts involved in RNA splicing/processing (**Figure 6B**), and splice isoforms play critical roles in synaptic plasticity and memory storage (Poplawski et al., 2016), we next assessed effects of SD and CFC on differentially-spliced mRNA isoforms using transcript-level analysis (Yi et al., 2018) (**Figure 6C**). On cytosolic ribosomes from pS6+ neurons, both the activity-dependent splice isoform of Homer1 scaffolding protein (*Homer1a*) and the highly-stable activity-dependent splice isoform of FosB (*ΔFosb*) were increased after CFC followed by *ad lib* sleep, but not after CFC in SD mice (**Figure 6C**). These isoforms were also increased as a function of SD alone, suggesting another mechanism by which SD could occlude changes to pS6+ neurons initiated by learning (**Figure 6C, Yellow**).

To validate and extend these findings, we harvested hippocampi from CFC and HC mice following 5 h of SD or *ad lib* sleep (i.e., a later time point with respect to learning). Using qPCR, we first quantified mRNA levels for splice isoforms of *Fosb* and *Homer1* in the cytosolic fraction of whole hippocampus (Input), Camk2a+ neurons, and pS6+ neurons. Similar to what was observed after 3 h of SD, 5 h of SD increased expression of both *Fosb* and its long-lasting splice isoform *ΔFosb*, regardless of prior CFC (**Figure 8A, left**). Compared with Input, *Fosb* and *ΔFosb* transcripts were relatively de-enriched in Camk2a+ neurons, but highly enriched in pS6+ neurons (consistent with neural activity regulating both S6 phosphorylation and *Fosb* and *ΔFosb* transcript abundance) (**Figure 8A, right**). To measure CFC-driven changes in these transcripts, we normalized *Fosb* and *ΔFosb* transcripts in CFC + Sleep or CFC + SD mice to that of the corresponding HC control group. In both pS6+ neurons and Input, *ΔFosb* increased following CFC, regardless of subsequent sleep or SD. In Camk2a+ neurons, however, both *Fosb* and *ΔFosb* transcripts increased in CFC + Sleep mice, but this increase was occluded in CFC + SD mice (**Figure 8A, bottom**).

**Figure 8.**
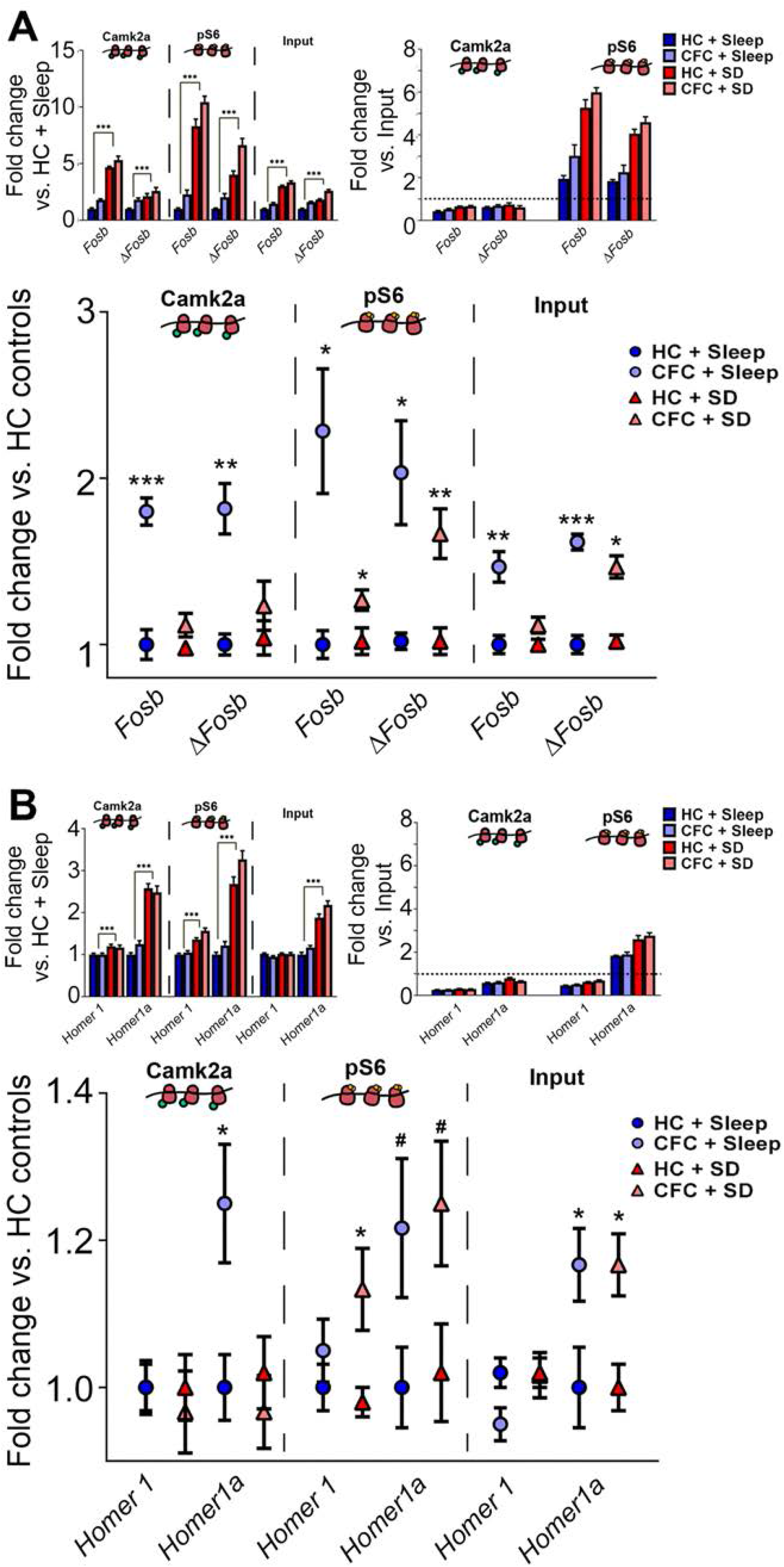
CFC-induced alterations in *Fosb* and *Homer1* splice variants are occluded by post-CFC SD in Camk2a+ neurons. **(A)** *Fosb* and *ΔFosb* expression for CFC and HC mice after 5 h of subsequent sleep or SD is shown for cytosolic fractions of the 3 cell populations. ***Top, left:*** Expression in the 4 conditions relative to values from HC + Sleep mice (Two-way ANOVA, CFC/SD, df = 18, *** p-value < 0.001). ***Top, right:*** Relative enrichment/de-enrichment for *Fosb* and *ΔFosb* in Camk2a+ and pS6+ neurons, relative to Input, for the 4 conditions. ***Bottom:*** Expression of Fosb and *ΔFosb* following CFC conditions relative to same-state (SD or Sleep) home-cage (HC) conditions (t-test, *n* = 5/group[HC], *n* = 6/group[CFC]) #, *, **, and *** indicate *p* < 0.1, *p* < 0.05, *p* < 0.01, and *p* < 0.001, respectively. **(B)** *Homer1* and *Homer1a* expression in cytosolic fractions in the 4 conditions, normalized as described in **(A)**.

We also used qPCR to quantify the relative expression *Homer1* and its splice variant *Homer1a* in cytosolic fractions after CFC and 5 h subsequent sleep or SD. *Homer1* itself was modestly affected by 5 h of SD, whereas the *Homer1a* transcript was dramatically elevated, consistent with earlier findings (Mackiewicz et al., 2008; Maret et al., 2008) (**Figure 8B, left**). *Homer1* was de-enriched in both Camk2a+ and pS6+ neurons relative to Input, while *Homer1a* was enriched only in pS6+ neurons (consistent with regulation by neuronal activity) (**Figure 8B, right**). Similar to *ΔFosb*, CFC increased *Homer1a* across all cell populations in mice allowed *ad lib* sleep, but this increase was occluded in Camk2a+ neurons by SD (**Figure 8C, bottom**).

We quantified additional cytosol-enriched transcripts for proteins with known functions in hippocampal plasticity and memory, to test the effects of CFC and 5 h subsequent sleep or SD (**Figure S4**). These included transcripts increased in our Desq2 analysis following SD alone, and either unaffected by CFC (*Cfos, Arc*), altered only in CFC + SD mice (*Atf3*), or altered only in CFC + Sleep mice (*1700016P03Rik).* We also visualized *Egr3* which was unaffected by SD but increased by CFC in both CFC + SD and CFC + Sleep groups. All transcripts except *Egr3* were altered by 5 h SD in Camk2a+ and pS6+ neuron populations. In pS6+ neurons, *Cfos* and the lncRNA transcript *1700016P03Rik* (Aten et al., 2016; Aten et al., 2018) remained significantly elevated as a function of learning 5 h following CFC; SD either fully or partially occluded these learning-associated changes (**Figure S4**). No significant CFC-induced changes in these transcripts were detectable on cytosolic ribosomes from Camk2a+ neurons at 5 h post-CFC.

Taken together, these data support the hypothesis that CFC-associated changes in activity-regulated transcripts at cytosolic ribosomes are likely occluded by subsequent SD. This effect, which is most pronounced for highly-active (putative engram) hippocampal neurons, constitutes a plausible mechanism for memory consolidation disruption by SD.

### SD Affects MB Ribosomal Transcripts Involved in Receptor-Mediated Signaling, Endoplasmic Reticulum Function, and Protein Synthesis

Fewer mRNAs were altered as a function of SD alone on MB ribosomes compared with cytosolic ribosomes (where most observed changes were driven by SD, rather than CFC) (**Figure 4, Figure 9A**). Changes in MB ribosomal transcripts due to SD were also dwarfed by more numerous changes to MB ribosome-associated transcripts following CFC. These changes differed substantially between Camk2a+ neurons, pS6+ neurons, and Input, thus canonical pathways represented by transcripts altered in the three populations by SD also differed. Critically, no canonical pathways were significantly enriched by SD-induced transcript changes in either Camk2a+ neurons or Input. On MB ribosomes from both Input and Camk2a+ cell populations, SD-altered transcripts included components of cellular pathways that were significantly affected in SD-regulated transcripts on cytosolic ribosomes (see **Figure 6B**), and with transcripts affected by SD in prior whole hippocampus transcriptome studies (Naidoo et al., 2005; Tudor et al., 2016; Vecsey et al., 2012). These included components of the AMPK (*Chrm5, Irs2, Pfkfb3, Ppm1f, Prkab2, Prkag2, Smarcd2*), IGF-1 (*Elk1, Rasd1*), IL-3 (*Crkl, Foxo1*), relaxin (*Gnaz, Pde4b, Smpdl3a*), and neuregulin (*Errfi1*) signaling pathways in Camk2+ neurons, and components of glucocorticoid receptor signaling (*Elk1, Gtf2e2, Prkab2, Prkag2, Rasd1, Smarcd2, Taf12, Fos, Krt77, Ptgs2, Tsc22d3*), unfolded protein response (*Hspa5, Pdia6*), and endoplasmic reticulum stress (Calr, *Xbp1*) pathways in both Camk2a+ neurons and Input. Critically, however, only 30 (6%) of the 511 mRNAs previously reported to be altered by SD in whole hippocampus (Vecsey et al., 2012) overlapped with SD-affected mRNAs in the MB fraction of whole hippocampus (i.e., Input; **Figure S2**). This suggests that even within these common identified cellular pathways, individual transcripts altered by SD in on MB ribosomes may differ substantially from those reported previously.

**Figure 9.**
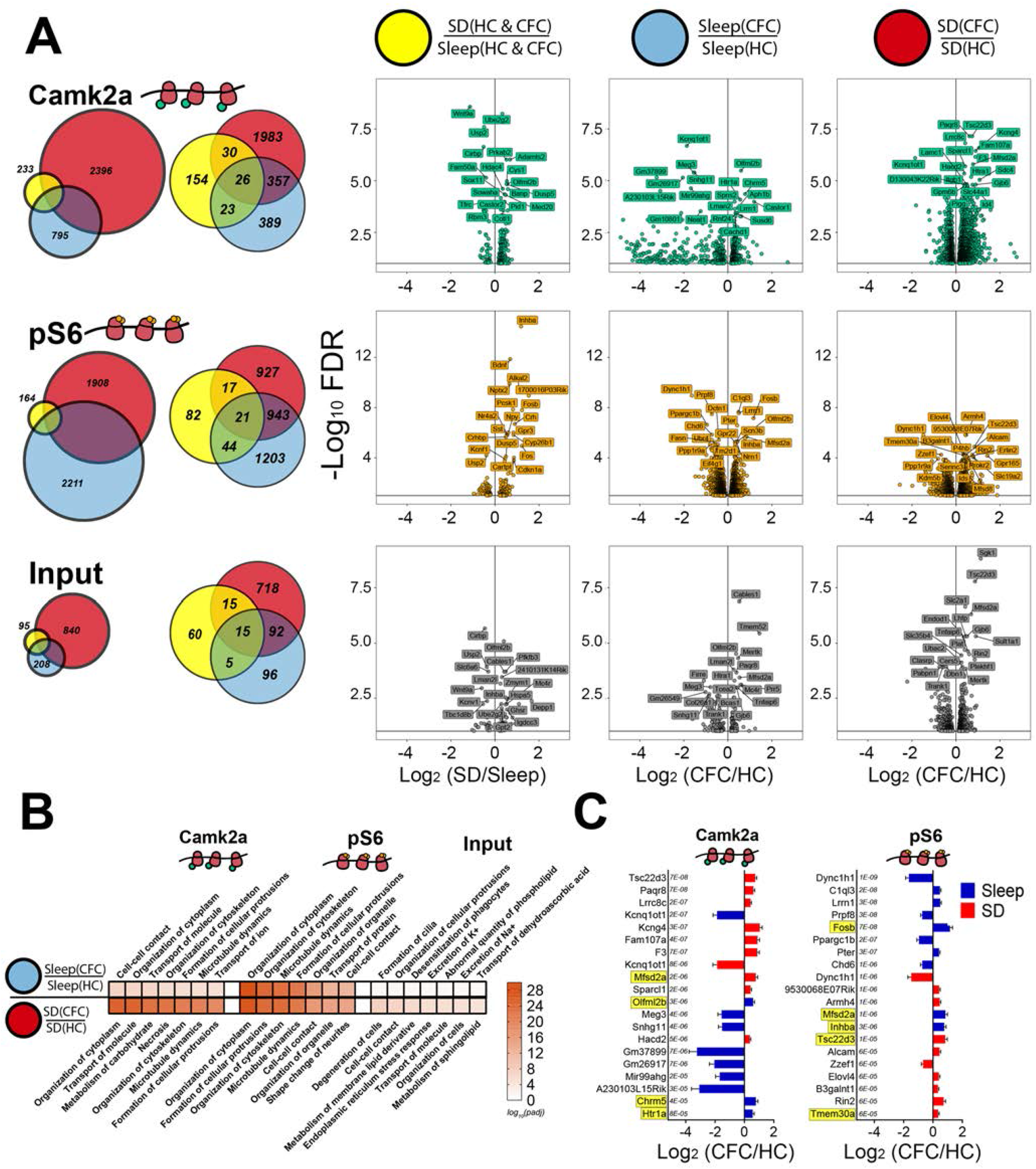
Transcripts altered by CFC on MB ribosomes encode regulators of neuronal morphology, intracellular trafficking, and lncRNAs. **(A) *Left*:** Proportional and overlapping Venn diagrams of transcripts significantly altered by SD, CFC + Sleep, and CFC + SD in MB fractions from Camk2a+ neurons, pS6+ neurons, and Input. ***Right:*** Volcano plots of Deseq2 results for transcripts measured in each condition. **(B)** The 7 most-significant molecular functions (ranked by *p*_adj_ value) for transcripts altered by CFC + Sleep (***top***) and CFC + SD (***bottom***) in Camk2a+ neurons, pS6+ neurons, and Input. **(C)**The 10 transcripts most significantly affected in CFC + Sleep (blue) and CFC + SD (red) conditions for Camk2a+ and pS6+ neurons, ranked by *p*_adj_ value. Transcripts that were also significantly altered as a function of SD alone are highlighted in yellow. Functional category analysis available in **Supplemental Table S10.**

In contrast to SD-driven changes in Camk2a+ neurons and Input, SD-altered transcripts from pS6+ neurons’ MB ribosomes significantly enriched for several canonical pathways. These included the neurotrophin/TRK (*Atf4, Bdnf, Fos, Ngf, Plcg1, Spry2*), corticotropin releasing hormone (*Crh, Vegfa*), ERK5 (*Il6st, Rasd1*), and EIF2 (*Eif2b3, Hspa5, Ptbp1, Rpl37a*) signalling pathways and the human embryonic stem cell pluripotency pathway (*Bmp2, Inhba, Wnt2*). Thus the major pathways affected by SD among MB ribosome-associated transcripts comprise receptor signalling pathways, protein synthesis regulation, and endoplasmic reticulum function.

### Learning-Related Changes in MB Ribosomal Transcripts Diverge Based on Subsequent Sleep or SD

In contrast to the sparsely expressed CFC-driven transcript changes observed on cytosolic ribosomes, the majority of transcript changes on MB ribosomes were driven by CFC (**Figure 4, Figure 9A**). Critically, learning-induced changes in MB-fraction transcripts diverged in all cell populations, depending on whether CFC was followed by 3 h of *ad lib* sleep or 3 h of SD. In contrast to the high degree of overlap between SD-driven and CFC-driven transcript changes in the cytosol, on MB ribosomes, mRNAs affected by SD showed significantly less proportional overlap (2-10%) with CFC-induced changes (**Figure 9A**). These data suggest that translational profiles of MB ribosomes are most selectively affected by prior learning, but that the specific mRNAs associated with MB ribosomes also vary dramatically as a function of post-learning sleep or SD.

We first characterized the cellular and molecular functions of MB ribosomal mRNAs altered as a function of CFC and subsequent sleep or SD. For Camk2a+ and pS6+ neurons, the most enriched functional categories largely overlapped, and represented similar molecular categories in CFC + Sleep and CFC + SD mice - including organization of cytoplasm, organization of cytoskeleton, microtubule dynamics, cell-cell contact, and formation of protrusions (**Figure 9B**). In contrast, few of these categories were enriched in Input MB fractions. There, the most enriched functional categories for transcripts altered by CFC + Sleep included excretion of sodium and potassium, formation of cilia, organization of cellular protrusions, desensitization of phagocytes, and abnormal quantity of phospholipid. Alterations in Input mRNAs following CFC + SD also enriched for functional categories not represented in neuronal MB fractions, including cell degeneration, metabolism of membrane lipid derivative, metabolism of sphingolipid, endoplasmic reticulum stress response. Together, these data suggest that CFC may alter similar membrane-associated cellular functions in Camk2a+ and pS6+ neuronal populations, regardless of subsequent sleep or SD. In contrast, CFC may have distinct and pronounced effects on membrane-associated functions in other hippocampal cell types (e.g., glia), and that these effects may diverge based on the animal’s sleep state.

To further characterize changes in MB-associated ribosomal transcripts following learning, we first compared significantly altered mRNAs (based on adjusted *p* value) in Camk2a+ and pS6+ neurons following CFC in sleeping and SD conditions (**Figure 9C**). At Camk2a+ neurons’ MB ribosomes, CFC + Sleep led to increased abundance for mRNAs encoding transmembrane receptors (*Chrm5, Htr1a*) and dramatically decreased abundance for multiple lncRNAs including *Kcnq1ot1, Meg3, Mir99ahg* and unannotated transcripts (e.g., *Gm37899, Gm26917*) (**Figure 9C**). Many other lncRNAs showed reduced abundance on MB ribosomes following CFC (including *Neat1, Malat1, Mirg*, and *Ftx).* With the exception of *Mirg* and *Ftx*, these lncRNAs were also significantly reduced following CFC + SD. CFC + SD led to the most significant transcript increases on MB ribosomes for *Lrrc8c* (encoding an acid sensing, volume-regulated anion channel), and anti-adhesive extracellular molecules (*Sparcl1/Hevin*), adhesion molecules (*F3/Contactin1*), transmembrane receptors (*Paqr8*), potassium modifiers (*Kcng4*), endoplasmic reticulum-tethered lipid synthesis molecules (*Hacd2*), and actin regulators (*Fam107a).*

In highly active (pS6+) neurons, *Dync1h1* was the most significantly altered transcript following CFC, and was dramatically reduced in both freely-sleeping and SD mice (**Figure 9C**). *Dync1h1* encodes the main retrograde motor protein in eukaryotic cells, supporting retrograde transport in axons and dendrites (Schiavo et al., 2013). To a lesser extent, its abundance was also significantly reduced on MB ribosomes from Camk2a+ neurons. *Dync1h1* was not reduced as a function of SD itself in either neuron population, suggesting that decreases in *Dync1h1* are specific to the post-learning condition.

We next constructed canonical pathway networks affected by CFC + Sleep or CFC + SD, to visualize the signaling and metabolic pathways differently altered in the two conditions. Canonical pathways are represented as hubs and connected through commonly transcript components. Here, hub sizes were weighted by their corresponding *p* value and shaded to indicate their z-score (blue indicating a decrease in the pathway following CFC, whereas red indicates an increase) (**Figure 9**). Network comparisons of MB ribosome-associated transcript changes in Camk2a+ neurons revealed overlapping hubs significant in both CFC + Sleep (**Figure 9, Top Left**) and CFC + SD conditions (**Figure 9, Top Right**). These hubs represented chondroitin sulfate degradation, unfolded protein response, notch signaling, phagosome maturation, sertoli cell junction signaling, and epithelial adherens signaling canonical pathways. However, the significance values for these pathway hubs (like the number of transcripts altered in Camk2a+ neurons after CFC) were markedly higher in SD mice relative to sleeping mice. For example, the overlap and centrality of sertoli cell junction signaling, phagosome maturation, and epithelial adherens junction signaling pathways suggest common transcripts altered in both freely-sleeping and SD mice after CFC. Upon closer inspection, while some tubulin transcripts were decreased following CFC in both Sleep and SD groups (*Tuba1a, Tuba1b*), a large number of tubulin-encoding mRNAs were decreased only after SD (*Tuba4a, Tubb2a, Tubb3, Tubb4a, Tubb4b, Tubg1*) (**Supplemental Table S10**). Similarly, while unfolded protein response-related transcripts were moderately elevated following CFC + Sleep (*Calr, Mbtps1, P4hb, Sel1l, Syvn1*), substantially more mRNAs associated with the unfolded protein response were increased after CFC + SD (*Amfr, Calr, Canx, Cd82, Cebpz, Dnajc3, Edem1, Eif2ak3, Hsp90b1, Hspa5, Mapk8, Mbtps1, Nfe2l2, Os9, P4hb, Sel1l, Syvn1, Ubxn4*, and *Xbp1).*

In Camk2a+ neurons, CFC + SD also altered the expression of MB ribosome-associated mRNAs linked to metabolic pathways that were unaffected in the CFC + Sleep group **(Figure 10A**). For example, CFC + SD increased abundance of transcripts related to lipid (triacylglycerol, phosphatidylglycerol, cdp-diacylglycerol) biosynthesis, including mRNAs encoding 1-acylglycerol-3-phosphate O-acyltransferases (*Agpat2, Agpat3, Agpat4*), ELOVL fatty acid elongases (*Elvol1, Elovl2, Elovl6*), and phospholipid phosphatases (*Plpp3*). CFC + SD also decreased the abundance of transcripts related to glucose metabolic pathways (glycolysis, gluconeogenesis, and TCA Cycle), including *Aldoa, Adloc, Eno1, Eno2, Gapdh, Gpi1, Pfkl*, and *Pkm* (**Supplemental Table S10**). Taken together with results shown in **Figure 9**, these data indicate that Sleep and SD lead to divergent changes in the bioenergetic genes present at membrane-bound Camk2a+ ribosomes following learning.

**Figure 10.**
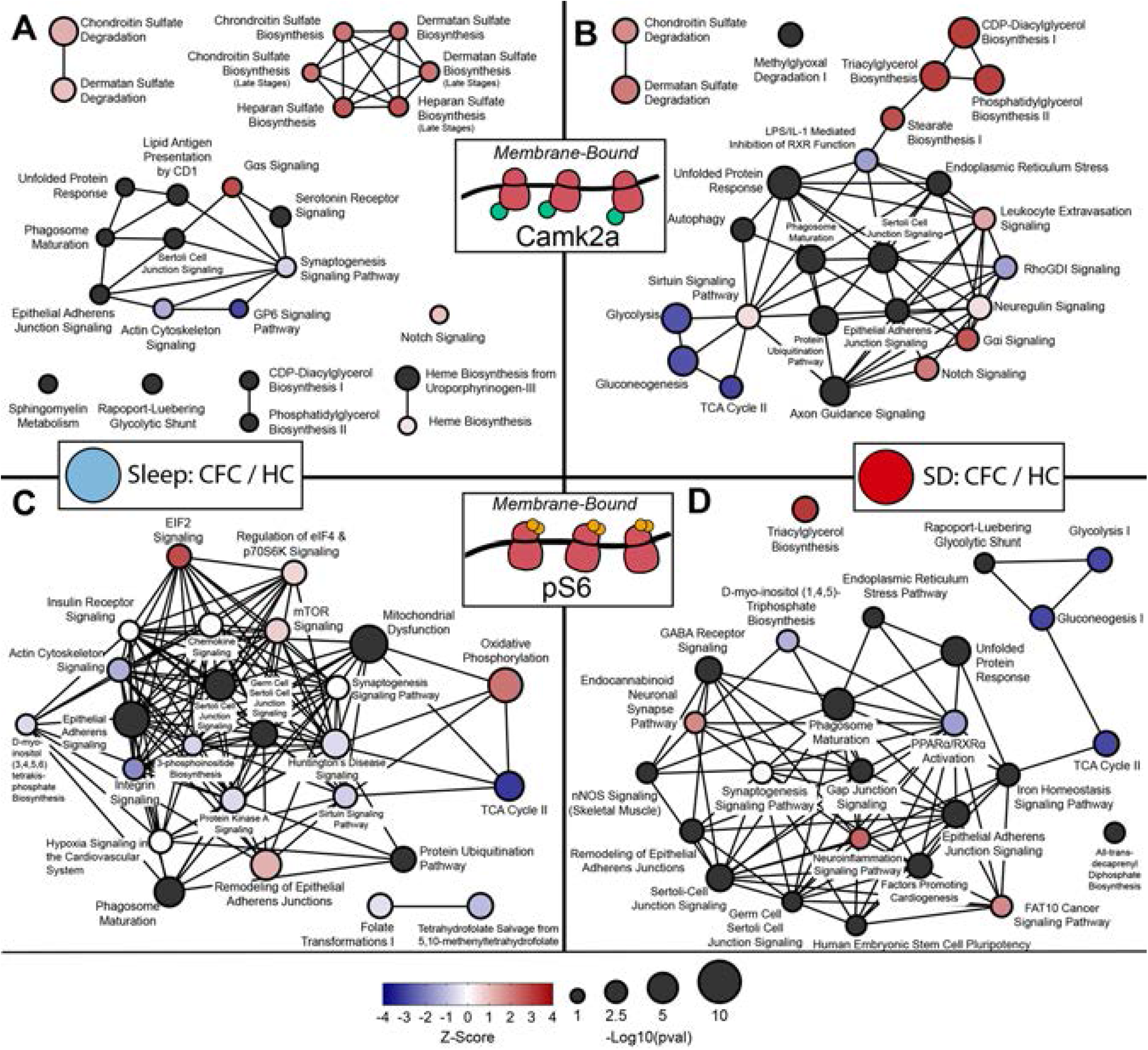
MB ribosomal transcript networks affected by CFC vary as a function of subsequent sleep or SD. Canonical pathway network analysis of transcripts altered on MB ribosomes from Camk2a+ (***top***) or pS6+ (***bottom***) neurons following CFC + Sleep (***left***) or CFC + SD (***right**).* Hub size and color denote *p*_adj_ value and z-score, respectively, in each condition, while connecting lines indicate commonly expressed genes between hubs. Canonical pathways are available in **Supplemental Table S10.**

We performed a similar canonical pathway network analysis on transcripts altered on MB ribosomes from pS6+ neurons following CFC (**Figure 10, Bottom**). Many of the same pathways altered by CFC in Camk2a+ neurons (in both Sleep and SD conditions) were also observed in pS6+ neurons - including sertoli cell junction signaling, epithelial adherens signaling, and phagosome maturation - suggesting some overlap. Pathways affected in the CFC + SD condition in Camk2a+ neurons included lipid and carbohydrate pathways affected in pS6+ neurons. Interestingly, in the CFC + Sleep condition (where CFM is being consolidated), there was an increased abundance of MB ribosome-associated transcripts representing protein translation regulatory pathways (eIF2, regulation of eIF4 & p70S6K, and mTOR signaling pathways). This change, critically, was not present in CFC + SD mice. In both sleeping and SD mice, MB ribosomal transcripts which decreased in abundance after CFC included eukaryotic initiation factors (*Eif3a, Eif3c, Eif3l, Eif4a1, Eif4g1, Eif4g3), mTOR*, and *Tsc1.* However, in mice allowed post-CFC sleep, transcripts related to the small ribosomal subunit were elevated in pS6+ neurons, including *Rps12, Rps14, Rps17, Rps19, Rps20, Rps21, Rps23, Rps24, Rps26, Rps28, Rps29, Rps6.* This suggests that following CFC, sleep may promote an increase in overall protein synthesis capacity, which occurs in the most active hippocampal neurons. The fact that these changes occur selectively on MB ribosomes suggest that this increased synthetic capacity may be cell compartment-specific.

## Discussion

Our present RNAseq results demonstrate not only that ribosome-associated transcripts are altered in the hippocampus as a function of 1) learning and/or 2) sleep vs. sleep loss, but also as a function of 3) the cell population being profiled, and 4) the subcellular location of the ribosomes. We find that the latter aspect (i.e., location of ribosomes within the cell) is a major contributor to the observed effects of learning and subsequent sleep or SD on hippocampal ribosome transcript profiles. Neuronal ribosomes have long been known to segregate by cell compartment, present either as “free-floating” (i.e., cytosolic) or MB complexes which are easily separated by centrifugation (Andrews and Tata, 1971). These populations are known to engage in compartmentalized translation of mRNAs. The advent of TRAP has yielded new insights into the specialized functions of ribosomes in different cellular compartments. Cytosolic ribosomes are known to process mRNAs encoding proteins with functions in the cytosolic compartment, including transcription factors and kinases. MB ribosomes translate mRNAs encoding secreted or integral membrane proteins. Available data, from non-neural cell types, suggest that the two translational environments are biochemically distinct and can be differentially regulated (for example, by cellular stress) (Reid and Nicchitta, 2015). Where ribosomes have been isolated from subcellular compartments in neurons (e.g. in Purkinje neurons) (Kratz et al., 2014), MB ribosome fractions have been shown to enrich for endoplasmic reticulum-associated ribosomes and for ribosomes in the dendritic compartment engaged in local translation. Our present findings reflect this, demonstrating that the transcript profiles of MB and cytosolic ribosomes among hippocampal neurons are highly distinctive (**Figure 1-3**, **Figure S1**).

Many forms of hippocampus-dependent memory are disrupted (in human subjects and animal models) by either pre- or post-learning sleep loss (Havekes and Abel, 2017; Krause et al., 2017; Puentes-Mestril and Aton, 2017; Rasch and Born, 2013). Indeed, sleep loss seems to disrupt plasticity mechanisms within the hippocampus more dramatically than in other brain areas (Delorme et al., 2019; Raven et al., 2019). The underlying mechanisms by which sleep loss leads to these changes (and disrupts memory mechanisms) have remained elusive. Transcriptome profiling of the effects of sleep loss alone on the hippocampus has indicated that SD increases expression of genes involved in transcriptional activation, and downregulates expression of genes involved in transcriptional repression, ubiquitination, and translation (Vecsey et al., 2012). While neither our cytosolic, MB, or Input hippocampal fractions showed a large degree of overlap with transcripts affected in prior studies (**Figure S2**), we do find that the same cellular pathways are affected in the cytosolic fraction (**Figure 6B**). Our present data add to this by demonstrating that in both whole hippocampus, and in either Camk2a+ or pS6+ hippocampal neurons, the majority of purely SD-induced changes to transcripts are present in the cytosolic fraction, and on cytosolic ribosomes (**Figure 4B, Figure 6**). While SD-driven mRNA changes also occur on MB ribosomes, these changes, in comparison, are relatively few in number (**Figure 4B, Figure 9**). Pathways affected by SD - across neuronal populations and both subcellular compartments - were those linked to regulation of transcription (consistent with prior findings) (Vecsey et al., 2012) and the AMPK, IL-3, IGF-1, and PDGF signaling pathways. Critically, AMPK (Chikahisa et al., 2009) and IGF-1 signaling (Chennaoui et al., 2014) have been implicated in homeostatic sleep responses (changes in sleep architecture of brain oscillations) following SD. Thus it is tempting to speculate that sleep loss could lead to subsequent changes in sleep brain dynamics through changes in intracellular signaling in neurons.

However, two unanswered questions are 1) which SD-associated changes in specific transcripts’ synthesis or translation provide a plausible mechanism to disrupt hippocampal memory consolidation, and 2) what cell types within the hippocampus are critically affected by SD following learning. This study aimed to address this in the context of a form of hippocampus-dependent memory consolidation (contextual fear memory; CFM) which is critically dependent on post-learning sleep. Work from our lab and others has shown that disruption of sleep within the first few hours following CFC is sufficient to disrupt CFM consolidation (Graves et al., 2003; Ognjanovski et al., 2018). While some of systems-level mechanisms occurring during post-CFC sleep have been implicated in the consolidation process (Boyce et al., 2016; Ognjanovski et al., 2018; Ognjanovski et al., 2014; Ognjanovski et al., 2017; Xia et al., 2017)), almost nothing is known about the cellular mechanisms mediating sleep (or SD) effects on CFM consolidation. We were surprised that very few transcript changes were induced on cytosolic ribosomes by CFC, in comparison with the large number of cytosolic ribosomal mRNAs affected by SD alone. However, of those transcripts altered by CFC, almost all 1) were similarly affected in either CFC + Sleep or CFC + SD conditions, and 2) were similarly altered by SD (**Figure 6C**). This makes sense in light of the fact that many cytosolic ribosomal mRNAs altered after CFC are transcribed or translated in an activity-dependent manner. We found that CFC increased the expression of activity-dependent genes in both total hippocampal mRNA (Input) and in the most activated neurons (pS6+) in both Sleep and SD mice. Importantly, post-CFC SD occluded learning-induced changes in cytosolic ribosomal transcripts present in excitatory Camk2a+ hippocampal neurons. These included increases in *Fosb* (and its highly stable isoform *ΔFosb*) and *Homer1* (and its short isoform *Homer1a*) (**Figure 8, Figure S4**). These transcript isoforms encode proteins that are critically linked to synaptic plasticity and memory (Clifton et al., 2019; Eagle et al., 2015). The timing with which SD occludes changes in their abundance (3-5 h following CFC) coincides a critical window for post-CFC sleep, essential for CFM consolidation (Graves et al., 2003; Ognjanovski et al., 2018). Thus this could represent a plausible mechanism for memory disruption by sleep loss.

In contrast to the relative paucity of transcripts altered on cytosolic ribosomes by prior learning, CFC affected a surprisingly large number of mRNAs on MB ribosomes (**Figure 4B, Figure 9A**). In general, CFC induced changes in MB ribosome-associated transcripts encoding proteins associated with neuronal structural remodeling - from cellular pathways involved in cytoskeletal remodeling, intracellular transport, and cell-cell interactions (**Figure 9B**). Some changes were also highly surprising and unexpected - for example, the significant reduction in ribosome-associated lncRNAs on MB ribosomes in Camk2a+ neurons after CFC (**Figure 9C**). Critically, the precise transcripts and (in some cases) the cellular pathways altered after CFC diverged dramatically based on whether learning was followed by sleep or SD (**Figures 9-10**). These differences provide a wealth of information with regard to potential mechanisms for SD-related disruption of CFM. For example, our present findings suggest that in non-neuronal cell types in the hippocampus, CFC induces a unique set of transcript changes (which are present in the MB fraction of Input, but are absent from MB fractions of neuronal ribosomes. Increased abundance of transcripts related to energy metabolism, particularly those encoding mitochondrial proteins, glucose transporters, and proteins related to glycogen metabolism are commonly observed following SD (Cirelli et al., 2004; Mackiewicz et al., 2007; Vecsey et al., 2012). Unlike previous reports, our data suggest that in Camk2a+ neurons, cellular metabolic/energetic pathways may be selectively disrupted when CFC is followed by SD, but not when CFC is followed by sleep (**Figures 9-10**). Thus, it may be that SD disrupts CFM consolidation by increasing the metabolic demands on the hippocampus.

In the most active (pS6+) neurons, CFC + Sleep leads to regulation of numerous pathways linked to protein synthesis regulation, including a widespread increase in MB ribosomal mRNAs encoding the translational machinery itself. This change is not seen when sleep is followed by SD. Thus it is tempting to speculate that in the neurons most active following memory encoding (putative “engram neurons”), long lasting changes to membrane associated protein synthesis may play a critical role in subsequent consolidation. Neuropharmacological studies have suggested that disruption of either cAMP signaling and protein synthesis in the hippocampus by SD may prevent memory consolidation (Abel et al., 1997; Bourtchouladze et al., 1998; Tudor et al., 2016; Vecsey et al., 2009). Our present data demonstrate that CFC may initiate changes to these pathways in specific hippocampal cell types, which are facilitated long-term by subsequent sleep.

Recently, TRAP has been used to characterize compartment-specific ribosomal transcripts of amygdala-projecting cortical axons during cued fear memory consolidation (Ostroff et al., 2019). However, the methods used in this study (as is true for most transcriptome and TRAP studies) would primarily report transcript changes associated with cytosolic, rather than MB ribosomes. Here we show that the vast majority of changes due to learning itself are expressed at MB ribosomes (**Figure 4B, Figure 9**) - with surprisingly few CFC-induced changes to cytosolic ribosome-associated mRNAs (**Figure 6**). Recent comparisons of hippocampal ribosome-associated and total mRNA abundance suggests that cytosolic and MB ribosome-associated mRNAs are distinctly regulated with regard to translation efficiency (Cho et al., 2015). Thus understanding the effects of both learning and subsequent sleep on structures like the hippocampus will require further investigation into their effects on translation happening at the membrane.

How universal are these sleep-dependent mechanisms for memory consolidation? While this question is presently unanswered, consolidation of various types of memory, across species, share common cellular substrates (Kandel et al., 2014), with post-learning mRNA translation being a vital element. Changes in the activity patterns of neurons and the activation of particular intracellular pathways during post-learning sleep share common features, across brain structures and species (Puentes-Mestril and Aton, 2017; Puentes-Mestril et al., 2019). Our present findings have illustrated a number of sleep-dependent post-learning cellular processes which affect pathways vital for learning and memory. Future studies will determine whether these processes underlie sleep-dependent memory consolidation events in the other brain circuits, following diverse forms of learning.

## Methods

### Mouse Husbandry, Handling, and Behavioral Procedures

All animal husbandry and experimental procedures were approved by the University of Michigan Institutional Animal Care and Use Committee (PHS Animal Welfare Assurance number D16-00072 [A3114-01]). For all studies, mice were maintained on a 12:12h light/dark cycle (lights on at 8 AM) with food and water provided *ad lib.* B6.Cg-Tg(Camk2a-cre)T29-1Stl/J mice (Jackson) were crossed to B6N.129-Rpl22^*tm1.1Psam*^/*J* mice (Jackson) to express HA-tagged Rpl22 protein in Camk2a+ neurons.

Mice were individually housed with beneficial enrichment for one week prior to experiments, and were habituated to handling (5 min/day) for five days prior to experiments. Mice were randomly assigned to one of four groups: HC + Sleep (*n* = 8), HC + SD (*n* = 7), CFC + Sleep (*n* = 8), CFC + SD (*n* = 7). Beginning at lights-on (8 AM), half of the mice underwent single-trial contextual fear conditioning (CFC) as described previously (Ognjanovski et al., 2018; Ognjanovski et al., 2014; Ognjanovski et al., 2017). Briefly, mice were placed in a novel conditioning chamber (Med Associates), and were allowed 2.5 min of free exploration time prior to delivery a 2-s, 0.75 mA foot shock through the chamber’s grid floor. After 3 min total in the chamber, mice were returned to their home cage (HC). As a control for the effects of learning, HC controls remained in their home cage during this time. HC + SD or CFC + SD mice were then kept awake continuously by gentle handling (SD; consisting of cage tapping, nest disturbance, and if necessary, stroking with a cotton-tipped applicator) over the next 3 h (for all RNA seq studies) or 5 h (for all qPCR experiments). HC + Sleep and CFC + Sleep mice were permitted *ad lib* sleep in their home cage for the same time interval.

### Translating Ribosome Affinity Purification (TRAP)

RiboTag TRAP was performed as previously described (Sanz et al., 2009) by indirect conjugation (Jiang et al., 2015), separating membrane-bound and free-floating ribosomes (Kratz et al., 2014). Briefly, following 3 h *ad lib* sleep or SD, mice were sacrificed with an overdose of pentobarbital (Euthasol). Brains were extracted and hippocampi dissected in ice cold dissection buffer (1x HBSS, 2.5 mM HEPES [pH 7.4], 4 mM NaHCO3, 35 mM glucose, 100ug/ml cycloheximide). Tissue was then transferred to glass dounce column containing 1 ml homogenization buffer (10 mM HEPES [pH 7.4], 150 mM KCl, 10 mM MgCl2, 2 mM DTT, 0.1 cOmplete™ Protease Inhibitor Cocktail [Sigma-Aldrich, 11836170001], 100 U/mL RNasin® Ribonuclease Inhibitors [Promega, N2111], and 100 μg/mL cycloheximide) and manually homogenized on ice. Homogenate was transferred to 1.5 ml LoBind tubes (Eppendorf) and centrifuged at 4°C at 1000 g for 10 min. The resulting supernatant (cytosolic fraction) was transferred to a new LoBind tube while the pellet (MB fraction) was resuspended in homogenization buffer. 10% NP40 was then added to the samples and incubated 5 min on ice, after which both MB and cytosolic fractions were centrifuged at 4°C at maximum speed for 10 min. The resulting supernatant from both MB and cytosolic fractions was then separated into Input (~50μL), Camk2a+ (~400μL), and pS6+ fractions (~500μL). For isolating ribosomes from Camk2a+ populations, fractions were incubated with 1:40 anti-HA antibody (Abcam, ab9110)(Shigeoka et al., 2018). To isolate ribosomes from highly active (pS6+) neurons fractions were incubated with 1:25 anti-pS6 244-247 (ThermoFisher 44-923G)(Knight et al., 2012). Antibody binding of the homogenate-antibody solution occurred over 1.5 h at 4°C with constant rotation.

For affinity purification, 200 μl/sample of Protein G Dynabeads (ThermoFisher, 10009D) were washed 3 times in 0.15M KCl IP buffer (10 mM HEPES [pH 7.4], 150 mM KCl, 10 mM MgCl2, 1% NP-40) and incubated in supplemented homogenization buffer (+10% NP-40). Following this step, supplemented buffer was removed, homogenate-antibody solution was added directly to the Dynabeads, and the solution was incubated for 1 h at 4°C with constant rotation. After incubation, the RNA-bound beads were washed four times in 900μL of 0.35M KCl (10mM HEPES [pH 7.4], 350 mM KCl, 10 mM MgCl2, 1% NP40, 2 mM DTT, 100 U/mL RNasin® Ribonuclease Inhibitors [Promega, N2111], and 100 μg/mL cycloheximide). During the final wash, beads were placed onto the magnet and moved to room temperature. After removing the supernatant, RNA was eluted by vortexing the beads vigorously in 350 μl RLT (Qiagen, 79216). Eluted RNA was purified using RNeasy Micro kit (Qiagen).

### Quantitative Real-Time PCR (qPCR)

For qPCR, RNA from each sample was converted into cDNA using the SuperScript IV Vilo Master Mix (Invitrogen 11756050). qPCR was performed on diluted cDNA that employed either Power SYBR Green PCR Mix (Invitrogen 4367659) or TaqMan Fast Advanced Master Mix (Invitrogen 4444557). For TRAP enrichment values, each sample was normalized to the geometric mean of *Pgk1* and *Gapdh* housekeeping transcripts and then normalized to the corresponding Input sample (TRAP Enrichment = 2^(ΔCt_target - ΔCt_housekeeping). Effects of SD were assessed by normalizing all groups’ expression to the HC + Sleep group. Effects of CFC were quantified by normalizing CFC + Sleep to HC + Sleep and normalizing CFC + SD to HC + SD. Primers for mRNAs quantified are listed below.

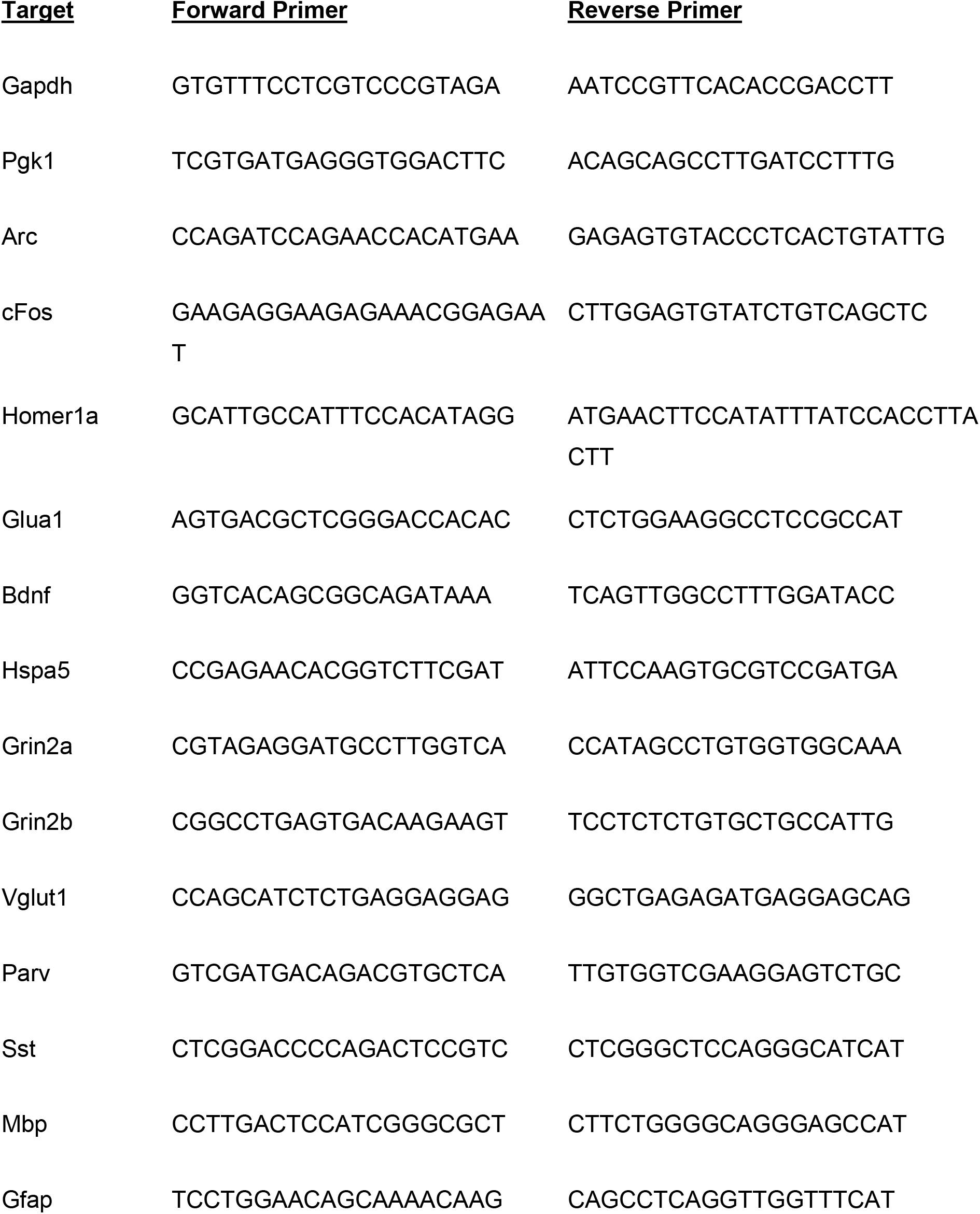

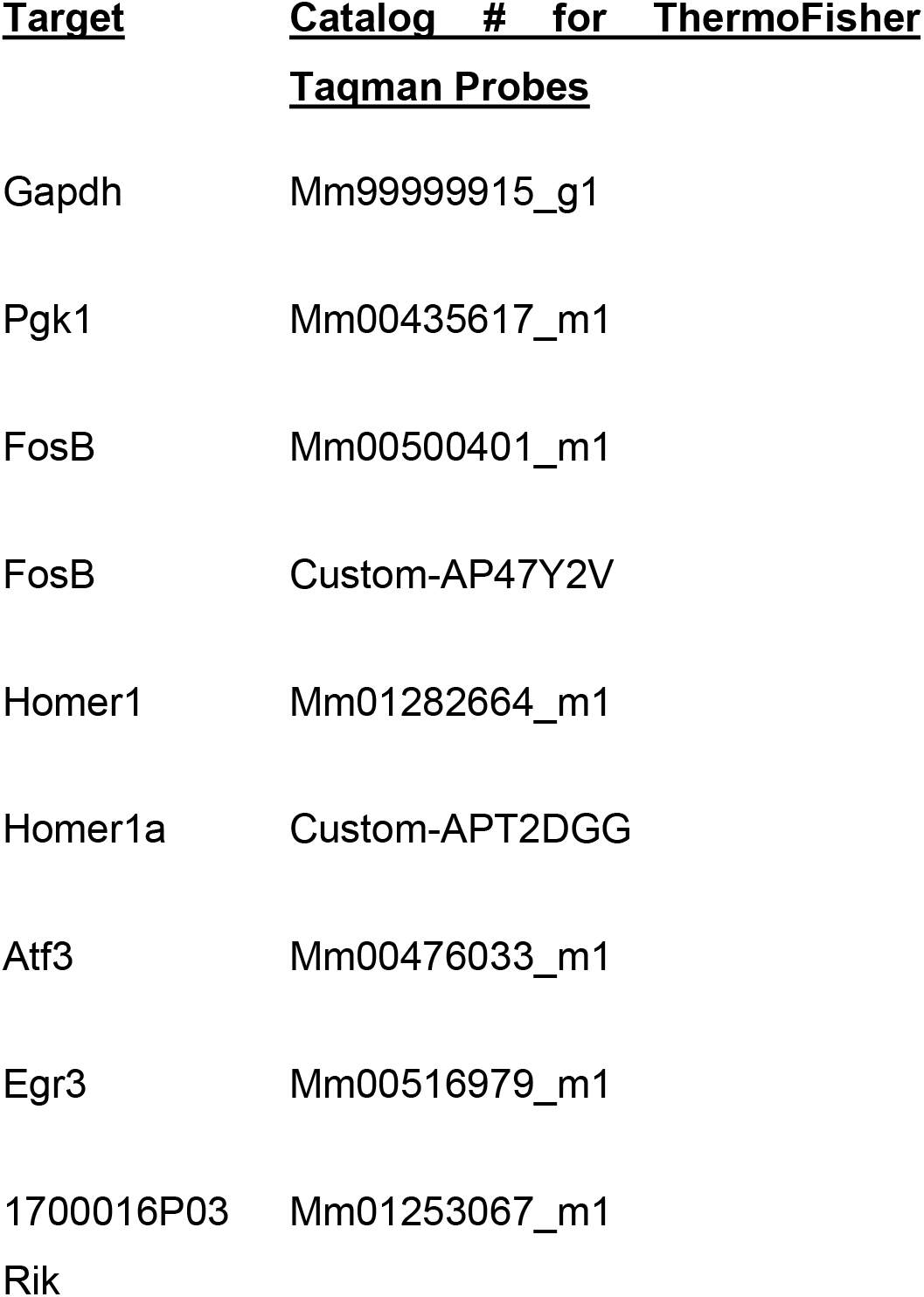

### RNA-Seq and Expression Analysis

RNA-Seq was carried out at the University of Michigan’s DNA Sequencing Core. Amplified cDNA libraries were prepared using Takara’s SMART-seq v4 Ultra Low Input RNA Kit (Takara 634888) and sequenced on Illumina’s NovaSeq 6000 platform. Sequencing reads (50 bp, paired end) were mapped to *Mus musculus* using Star v2.6.1a and quality checked with Multiqc (v1.6a0). Reads mapped to unique transcripts were counted with featureCounts (Liao et al., 2013).

Differential expression analyses were run with Deseq2 (Love et al., 2014). Analyses were run with an initial filtering step (removing rows with < 10 counts) and with betaPrior = False. To test differences between subcellular fractions within their respective cell population, the design of the GLM was set to compare differences between supernatant and pellet-enriched transcripts. Camk2a+, pS6+, and Input samples were analyzed separately (e.g., Camk2a+[supernatant/pellet]). To quantify effects of SD and CFC on expression, the design was switched and each cell population and subcellular fraction was analyzed separately. The same two-factor design was used to analyze the effects of an animal’s state (Sleep or SD) and learning (CFC or HC) on RNA expression. The design compared the effects of SD alone by combining HC and CFC animals. In contrast, the effect of learning (CFC) was assessed separately in CFC + Sleep and CFC + SD mice.

To characterize the differences between the effects of SD and CFC, significantly altered transcripts were analyzed using Ingenuity’s Pathway Analysis (IPA). GO analyses were performed in IPA and DAVID’s Functional Annotation tool. For subcellular fraction comparisons, 2000 of the top cytosolic (Log2FC > 0) and MB (Log2FC < 0) enriched transcripts (ranked by adjusted *p* values) were run through IPA’s Canonical Pathways analysis. To characterize differences in common metabolic pathways between cytosolic and MB fractions, hierarchical clustering was used to visualize the most differentially-expressed transcripts. Since signaling pathways were less overlapping between the MB and cytosolic fraction, they were ranked by enrichment *p* values. Those transcripts were then run through DAVID’s Functional Annotation tool, selecting for cellular composition to describe the cellular compartment the corresponding protein relates to. Data were plotted in Fragments Per Million (FPM) and their correlation value (R) calculated in the ViDger R package (McDermaid et al., 2019).

### Immunohistochemistry

To characterize HA and pS6 expression in the hippocampus, experimentally naive animals were sacrificed and perfused with 1xPBS followed by 4% paraformaldehyde. 50μM coronal sections containing dorsal hippocampus were blocked in normal goat serum for 2 h and incubated overnight using a biotin conjugated anti-HA (Biolegend 901505, 1:500), anti-pS6 244-247 (ThermoFisher 44-923G, 1:500), and anti-parvalbumin (Synaptic Systems 195 004, 1:500) antibodies. Sections were then incubated with secondary antibodies - Streptavidin-Alexa Fluor® 647 (Biolegend 405237), Fluorescein (FITC) Goat Anti-Rabbit IgG (H+L) (Jackson 111-095-003), and Alexa Fluor® 555 Goat Anti-Guinea pig IgG H&L (Abcam ab150186). Immunostained sections were coverslipped in ProLong Gold Antifade Reagent (ThermoFisher, P36930) for imaging with a Leica SP5 laser scanning confocal microscope.

## Acknowledgements

The authors are grateful to members of the Aton lab, and to Drs. Natalie Tronson and Ryan Mills for helpful feedback on this manuscript. This work was supported by research grants from the NIH (DP2 MH 104119) and the Human Frontiers Science Program (N023241-00_RG105) to SJA. We acknowledge support from the Bioinformatics Core of the University of Michigan Medical School’s Biomedical Research Core Facilities

