## Supplementary Figure legends for "Learning and Sleep Have Divergent Effects on Cytosolic and Membrane-Associated Ribosomal mRNA Profiles in Hippocampal Neurons"

### Supplementary Figure Legends and Tables.

**Figure S1. Transcripts enriched in cytosolic and MB fractions from Input (whole hippocampus). (A)** Volcano plot of transcripts significantly enriched in pellet (red) and supernatant (blue) cell fractions. Of the 27,773 transcripts detected, and 8,310 (30%) showed significant enrichment in the supernatant (cytosolic) fraction, 9,285 (33%) showed enrichment in the pellet (MB) fraction. Complete transcript list in **Supplemental Table S3**. **(B)** Top 10 cellular component localizations (from DAVID) of the 2000 transcripts which were most significantly enriched (based on $p_{adj}$ value) in either pellet (MB) or supernatant (cytosolic) fractions. **(C)** Top 20 most-enriched signaling and metabolic pathways represented by the 2000 most-enriched transcripts in Input cytosolic or MB fractions. **(D)** Log2FC values indicating enrichment of transcripts from the ubiquitin signaling pathway in the cytosolic (red) or MB (blue) fractions. Functional category analysis available in **Supplemental Table S3**

**Figure S2. Overlap of SD-altered transcripts with previously-characterized SD-altered mRNAs.** Venn diagrams indicate degree of overlap for transcripts altered by SD in the present study and those previously reported for whole hippocampus following SD (Vecsey et al., 2012).

**Figure S3. Input CREB upstream regulator analysis.** Networks of Creb1 transcriptional targets altered by SD in the cytosolic fraction taken from whole hippocampal homogenate (Input). Color of arrows indicates predicted regulation by Creb1 following SD whereas the color of the gene symbol indicates $Log_2$ FC in expression following SD. Orange arrows pointing to red symbols indicates genes increased by Creb1 that are also increased by SD while blue arrows connote transcripts predicted to be repressed by Creb1 which are also repressed by SD. Yellow arrows encompass all SD-related changes that do not match predicted regulation by Creb1; grey indicates undetermined effects of Creb1 on transcript levels. CREB network analyses are available in **Supplemental Figure 3**.

**Figure S4. CFC-induced alterations in activity-dependent transcripts in Sleep and SD mice.** Expression of activity-regulated transcripts *Cfos*, *Atf3*, *Arc*, and *Erg1*) and lncRNA *1700016P03Rik* for CFC and HC mice after 5 h of subsequent sleep or SD is shown for cytosolic fractions of Camk2a+ and pS6+ neurons. ***Top:*** Expression in the 4 conditions relative to values from HC + Sleep mice (Two-way ANOVA, CFC/SD, df = 18) * , ** , and *** indicate $p < 0.05$, $p <$

0.01, and $p < 0.001$, respectively. ***Bottom:*** Expression for the CFC conditions relative to same-state (SD of Sleep) HC conditions.

**Figure S5. Overview of data in Supplementary Tables.**

**Table S1 - Transcripts enriched in cytosolic or MB ribosomal fractions of Camk2a+ neurons.**

**Table S2 - Transcripts enriched in cytosolic or MB ribosomal fractions of pS6+ neurons.**

**Table S3 - Transcripts enriched in cytosolic or MB ribosomal fractions of Input.**

**Table S4 - Transcripts altered by SD and CFC in Camk2a+ neurons.**

**Table S5 - Transcripts altered by SD and CFC in pS6+ neurons.**

**Table S6 - Transcripts altered by SD and CFC in Input.**

**Table S7 - Venn diagram values.**

**Table S8 - Canonical/molecular functions altered by SD (cytosolic fractions).**

**Table S9 - CREB upstream regulator analysis (cytosolic fractions).**

**Table S10 - Canonical/molecular functions altered by SD and CFC (MB fractions).**
