## Supplementary figures and images for "Learning and Sleep Have Divergent Effects on Cytosolic and Membrane-Associated Ribosomal mRNA Profiles in Hippocampal Neurons"

### Figure S1

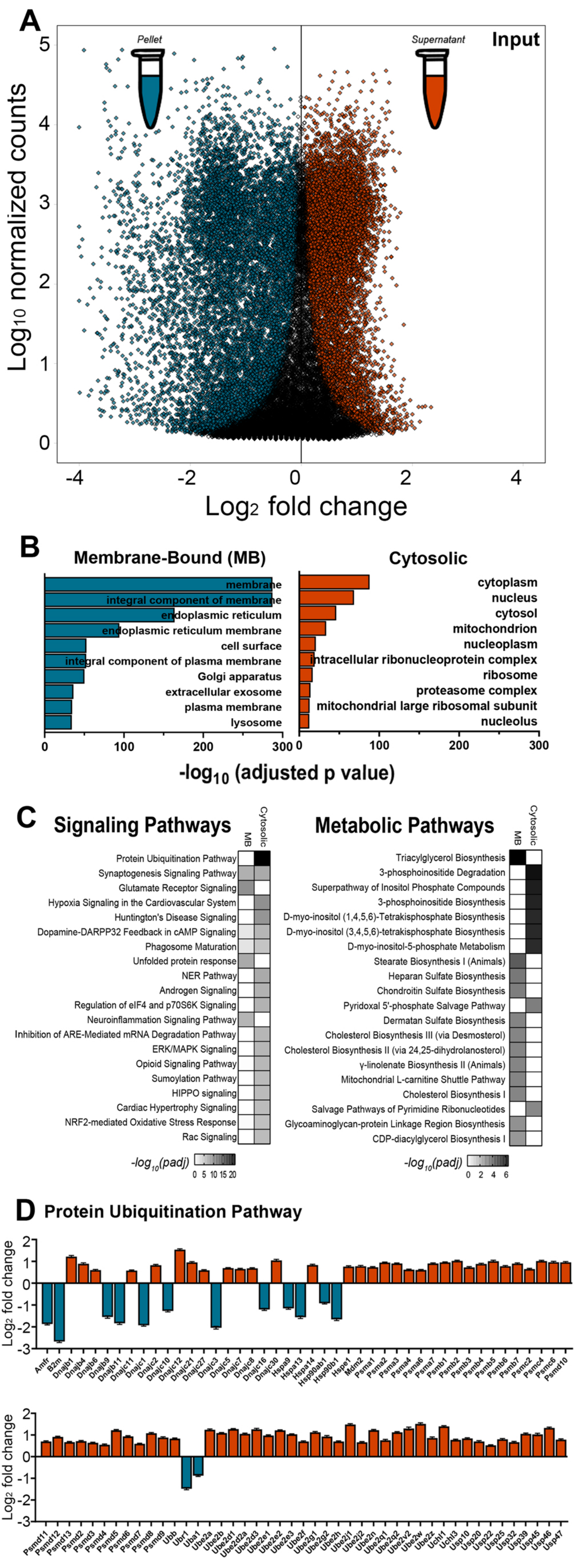

### Figure S2

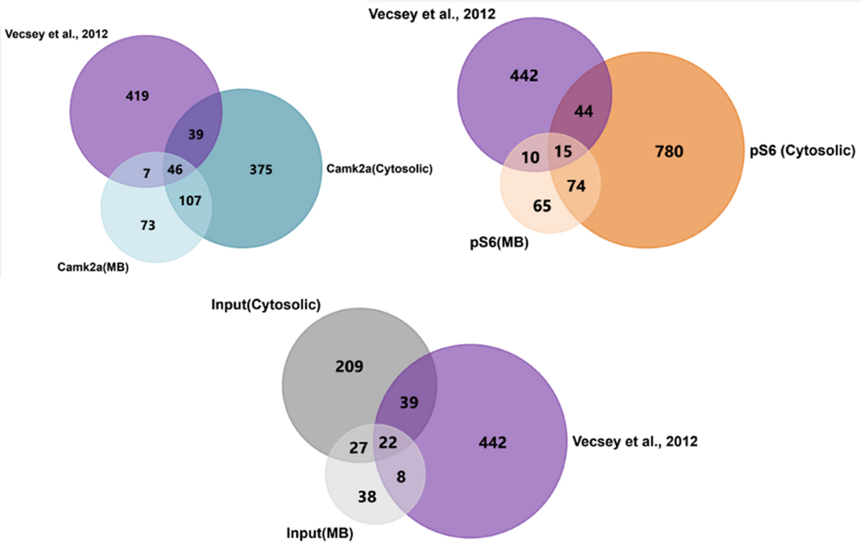

### Figure S3

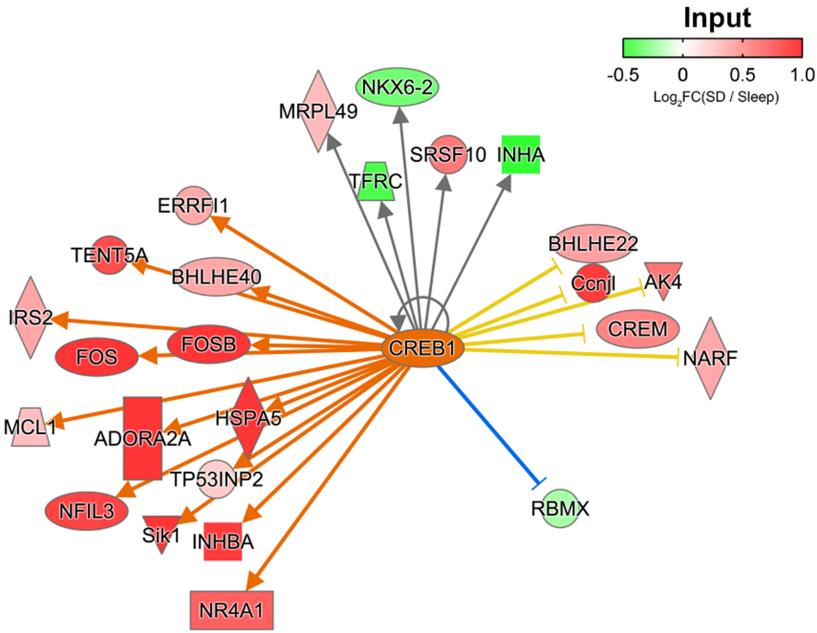

### Figure S4

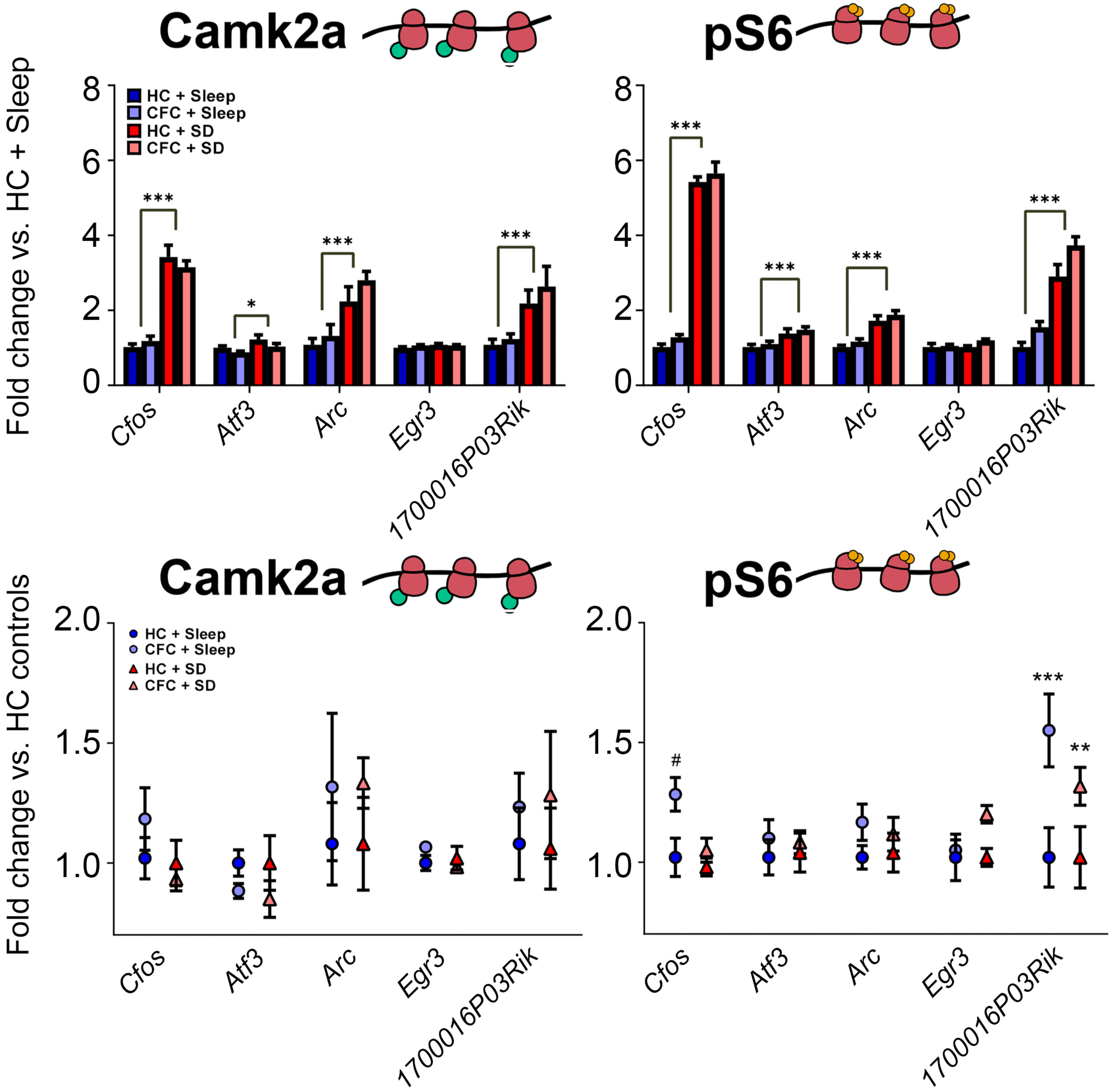
