## Supplementary material for "Learning and Sleep Have Divergent Effects on Cytosolic and Membrane-Associated Ribosomal mRNA Profiles in Hippocampal Neurons": Figure S5

### Overview of Supplemental Tables

#### Supernatant vs. Pellet Comparisons

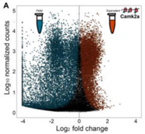


**Camk2a+** Supplemental Table 1

**pS6+** Supplemental Table 2

**Input+** Supplemental Table 3

#### SD & CFC Comparisons

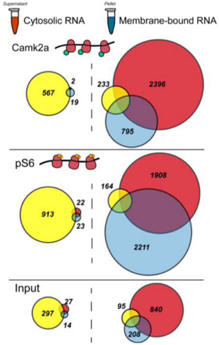


**Camk2a+** Supplemental Table 4

**pS6+** Supplemental Table 5

**Input+** Supplemental Table 6

Venn Overlap Transcripts (ALL), **Supplemental Table 7**

Canonical/Molecular Functions (Cytosolic), **Supplemental Table 8**

CREB UR Analysis (Cytosolic), **Supplemental Table 9**

Canonical/Molecular Functions (MB), **Supplemental Table 10**
